# Identity-by-descent detection across 487,409 British samples reveals fine-scale population structure, evolutionary history, and trait associations

**DOI:** 10.1101/2020.04.20.029819

**Authors:** Juba Nait Saada, Georgios Kalantzis, Derek Shyr, Martin Robinson, Alexander Gusev, Pier Francesco Palamara

## Abstract

Detection of Identical-By-Descent (IBD) segments provides a fundamental measure of genetic relatedness and plays a key role in a wide range of genomic analyses. We developed a new method, called FastSMC, that enables accurate biobank-scale detection of IBD segments transmitted by common ancestors living up to several hundreds of generations in the past. FastSMC combines a fast heuristic search for IBD segments with accurate coalescent-based likelihood calculations and enables estimating the age of common ancestors transmitting IBD regions. We applied FastSMC to 487,409 phased samples from the UK Biobank and detected the presence of ∼214 billion IBD segments transmitted by shared ancestors within the past 1,500 years. We quantified time-dependent shared ancestry within and across 120 postcodes, obtaining a fine-grained picture of genetic relatedness within the past two millennia in the UK. Sharing of common ancestors strongly correlates with geographic distance, enabling the localization of a sample’s birth coordinates from genomic data. We sought evidence of recent positive selection by identifying loci with unusually strong shared ancestry within recent millennia and we detected 12 genome-wide significant signals, including 7 novel loci. We found IBD sharing to be highly predictive of the sharing of ultra-rare variants in exome sequencing samples from the UK Biobank. Focusing on loss-of-function variation discovered using exome sequencing, we devised an IBD-based association test and detected 29 associations with 7 blood-related traits, 20 of which were not detected in the exome sequencing study. These results underscore the importance of modelling distant relatedness to reveal subtle population structure, recent evolutionary history, and rare pathogenic variation.

## Introduction

Large-scale genomic collection, through efforts like the NIH All of Us research program (1), the UK BioBank (2), and Genomics England (3), has yielded datasets of hundreds of thousands of individuals and is expected to grow to millions in the coming years. Utilizing such datasets to understand disease and health outcomes requires understanding the fine-scale genetic relationships between individuals. These relationships can be characterized using short segments (less than 10 centimorgans [cM] in length) that are inherited identical by descent (IBD) from a common ancestor between purportedly “unrelated” pairs of individuals in a dataset (4). Accurate detection of shared IBD segments has a number of downstream applications, which include reconstructing the fine-scale demographic history of a population (5–8), detecting signatures of recent adaptation (9, 10), discovering phenotypic association (11, 12), estimating haplotype phase (4, 13, 14), and imputing missing genotype data (15, 16), a key step in genome-wide association studies (GWAS) (17). Detection of IBD segments in millions of individuals within modern biobanks poses a number of computational challenges. Although several IBD detection methods have been published (18–20), few scale to analyses comprising more than several thousand samples. However, scalable methods that do exist trade modeling accuracy for computational speed. As a result, current IBD detection algorithms are either scalable but heuristic, solely relying on genotypic similarity to detect shared ancestry and not providing calibrated estimates of uncertainty, or too slow to be applied to modern biobanks. Here, we introduce a new IBD detection algorithm, called *fast sequentially Markovian coalescent* (FastSMC), which is both fast, enabling IBD analysis of modern biobank datasets, and accurate, relying on coalescence modeling to detect short IBD segments (down to 0.1 cM). FastSMC quantifies uncertainty and estimates the time to most recent common ancestor (TMRCA) for individuals that share IBD segments. It does so by efficiently leveraging information provided by allele sharing, genotype frequencies, and demographic history, which does not require computing shared haplotype frequencies and results in a cost-effective boost in accuracy.

We used extensive coalescent simulation to verify the scalability, accuracy, and robustness of the FastSMC algorithm in detecting IBD sharing within recent millennia. We then leveraged the speed and accuracy of FastSMC to analyze IBD sharing in 487,409 phased individuals from the UK Biobank dataset, identifying and characterizing ∼214 billion IBD segments transmitted by shared ancestors within the past 50 generations. This enabled us to reconstruct a fine-grained picture of time-dependent genomic relatedness in the UK. Analysis of the distribution of recent sharing within specific genomic regions revealed evidence of recent positive selection at 12 loci, 7 of which have not been previously reported. We found the sharing of IBD to be highly correlated with geographic distance and the sharing of rare variants. This gave us the opportunity to detect 20 novel associations to genomic loci harboring loss-of-function variants with 7 blood-related phenotypes.

## Results

### Overview of the FastSMC method

The algorithm we developed, called FastSMC, detects IBD segments using a two-step procedure. In the first step (identification), FastSMC uses genotype hashing to rapidly identify IBD candidate segments, which enables us to scale to very large datasets. In the second step (verification), each candidate segment is tested using a coalescent hidden Markov model (HMM), which enables us to improve accuracy, compute the posterior probability that the segment is IBD (*IBD quality score*), and provide an estimate for the TMRCA in the genomic region. The identification step leverages the new GERMLINE2 algorithm, which improves over GERMLINE’s (19) speed and memory requirements and thus enables us to very efficiently detect IBD candidate regions in millions of genotyped samples. GERMLINE2 utilizes hash functions to identify pairs of individuals whose genomes are identical in small genomic regions. The presence of these short identical segments triggers a local search for longer segments that are likely to reflect recent TMRCA and thus IBD sharing in the region. Although the original GERMLINE algorithm utilizes a similar strategy, GERMLINE2 offers two key improvements, which result in faster computation and lower memory consumption. First, the GERMLINE algorithm can become inefficient in regions where certain short haplotypes can be extremely common in the population (e.g. due to high linkage disequilibrium), which results in hash collisions across a large fraction of samples, effectively reverting back to a nearly all-pairs analysis and monopolizing computation time. GERMLINE2 avoids this issue by introducing recursive hash tables, which require haplotypes to be sufficiently diverse before they are explored for pairwise analysis and significantly decrease downstream computation, as shown in Supplementary Fig. 1. Second, the GERMLINE local search (extension) step requires storing the entire genotype dataset in memory, which is prohibitive for biobank-scale analyses. Instead, GERMLINE2 uses an *on-line* strategy, reading a polymorphic site at a time without storing complete genotype information in memory, which enables scaling this analysis to millions of individuals. The identification step efficiently finds genomic regions where the genotypes of a pair of individuals are the same, thus being “identical-by-state” (IBS). While long IBS regions are often co-inherited from recent common ancestors, thus being IBD, this need not always be the case (21). In its verification step, FastSMC thus leverages coalescence modeling to filter out candidate segments that are IBS, but not IBD. To achieve this, FastSMC analyzes every detected candidate segment using the ASMC algorithm (22), a recently proposed coalescent-based HMM that builds on recent advances in population genetics inference (23–26) to enable efficient estimation of the posterior of the TMRCA for a pair of individuals at each site along the genome. A key advantage of the ASMC algorithm over previous coalescent-based models is that it enables estimating TMRCA in SNP array data in addition to sequencing data. FastSMC can thus be tuned to be applied to both types of data. FastSMC produces a list of pairwise IBD segments with each segment associated to an *IBD quality score* - i.e the average probability of the TMRCA being between present time and the user-specified time threshold - and an age estimate - i.e the average maximum-a-posteriori (MAP) TMRCA along the segment. More details are described in the Methods. The FastSMC software implements both the GERMLINE2 (identification) and ASMC (validation) algorithms, and is freely available (see URLs).

Throughout this work, we define a genomic site to be shared IBD by a pair of phased haploid individuals if their TMRCA at the site is lower than a specified time threshold (e.g. 50 generations). This is a natural definition for IBD sharing, as it is closely related to several other quantities that are of interest in downstream analyses, such as genealogical relatedness or the probability of sharing rare genomic variants. We note, however, that a number of other definitions can be found in the literature (21). This is often due to the fact that current IBD detection algorithms cannot effectively estimate the TMRCA of a putative IBD segment. Downstream analyses of shared segments (e.g. (5, 8)) thus often resort to using the length of detected segments as a proxy for its age, since a segment’s length is expected to be inversely proportional to its TMRCA (see Methods).

### Comparison to existing methods

We measured FastSMC’s accuracy using extensive realistic coalescent simulations that mimic data from the UK Biobank (2) (see Methods for details on the simulated scenarios). We measured accuracy using the area under the precision-recall curve (auPRC), where precision represents the fraction of identified sites that are indeed IBD (following the TMRCA-based definition of IBD), and recall represents the fraction of true IBD sites that are successfully identified. We benchmarked IBD detection for FastSMC in addition to three other widely used or recently published IBD detection methods: GERMLINE (19), RefinedIBD (18), and RaPID (20). Parameters for all methods were optimized to maximize accuracy and evaluated on the detection of IBD segments within the past 25, 50, 100, 150, or 200 generations on simulated populations with a European ancestry (see Methods, Supplementary Table 1). We found that FastSMC outperforms the accuracy of all the other methods at all time scales (Fig. 1 **A, C**, Supplementary Fig. 2, Supplementary Table 2). As expected, model-based algorithms such as FastSMC and RefinedIBD tend to achieve better results in detecting older (shorter) IBD segments than genotype-matching methods, which cannot reliably exclude short segments where genotypes are identical (IBS) but not IBD (Supplementary Table 2). FastSMC relies on the ASMC algorithm in its validation step, which was shown to be robust to the use of an inaccurate recombination rate map or violations of assumptions on allele frequencies in SNP ascertainment (22). We thus expect it to be similarly robust to several types of model misspecification. In particular, we tested the effects of using a misspecified demographic model on FastSMC’s accuracy, and observed that while this results in biased estimates of segment age (Supplementary Fig. 3), a wrong demographic model does not affect auPRC accuracy (Supplementary Table 3).

**Fig. 1.**
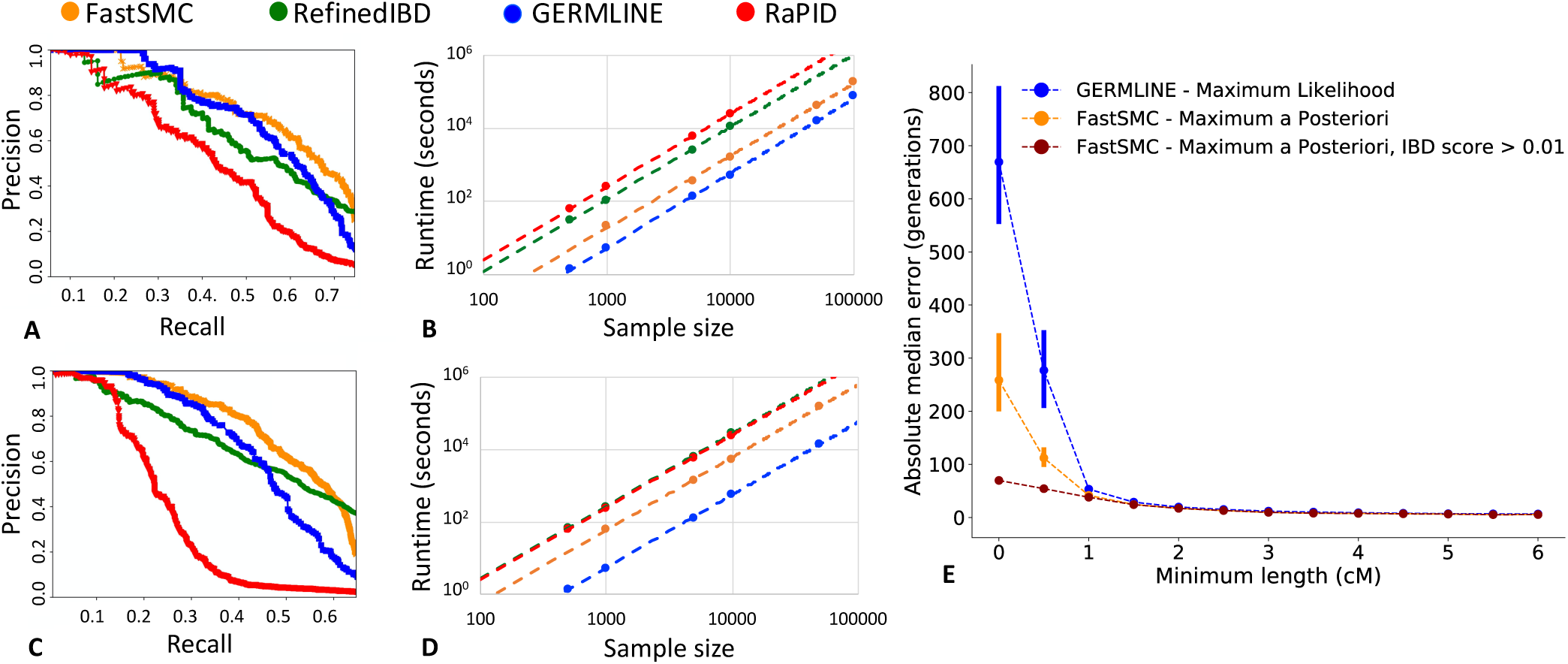
FastSMC in coalescent simulations. **A (respectively C).** Precision-recall curve randomly sampled from 10 realistic European simulated datasets with 300 haploid samples for IBD segments detection within the past 50 generations (respectively 100 generations), within the recall range where all methods are able to provide predictions. **B (respectively D).** Running time (CPU seconds) using chromosome 20 of the UK Biobank for IBD segments detection within the past 50 generations (respectively 100 generations). Only one thread was used for each method and running time trend lines in logarithmic scale are shown, reflecting differences in the quadratic components of each algorithm. Parameters for all methods were optimized to maximize accuracy and used for both accuracy and running time benchmarking (details in Methods). **E.** Absolute median error of the maximum likelihood age estimate of GERMLINE in blue (based on segments length only) and the MAP age estimate of FastSMC (no filtering on the IBD quality score in orange, and a minimum IBD quality score of 0.01 in dark red). Only IBD segments longer than the minimum length represented on the X-axis were considered. We ran both algorithms with the same minimum length of 0.001 cM and other parameters from grid search results for a time threshold of 50 generations (see Methods). Error bars correspond to standard error over 10 simulations.

Next, we evaluated the computational efficiency of FastSMC and other methods using phased data from chromosome 20 of the UK BioBank (Fig. 1 **B, D** and Supplementary Fig. 4). The entire cohort of 487,409 samples (across 7,913 SNPs) was randomly downsampled into smaller batches. As expected, the improvement in accuracy achieved leveraging FastSMC’s validation step leads to slightly increased computing time compared to only using GERMLINE, the most scalable method. FastSMC is faster than RefinedIBD, the closest method in terms of accuracy for short segments. For instance, detecting IBD segments within the past 50 generations on 10,000 samples takes 27 minutes for FastSMC, 9 minutes for GERMLINE, 3 hours and 17 minutes for RefinedIBD, and 6 hours and 58 minutes for RaPID. We finally assessed the memory cost of FastSMC and other methods (Supplementary Fig. 5). FastSMC does not store the genotype hashing or IBD segments, resulting in a very low memory footprint, whereas the memory requirements of other methods become prohibitive for large sample sizes such as those required to analyze the entire UK Biobank cohort. For example, analyzing chromosome 20 for a time threshold of 50 generations and 10,000 random diploid individuals from the UK Biobank dataset, FastSMC requires 1.4GB of RAM compared to 3.8GB for GERMLINE, 11.5GB for RefinedIBD and 62.9GB for RaPID.

Downstream analysis of IBD sharing such as demographic inference or the study of natural selection often involves estimating the age of IBD segments. Because current approaches do not explicitly model the TMRCA between IBD individuals, segment age is typically estimated through the length of the IBD segment (see Methods). FastSMC, on the other hand, explicitly models TMRCA across individuals, leveraging additional prior information (such as demography and allele frequencies) to produce an improved estimate of IBD segment age. We found FastSMC’s segment age estimates to be more accurate than a length-based estimator (Supplementary Fig. 6), with significant gains for short segments as a result of the additional modeling in the validation step of the algorithm (e.g. the median error from FastSMC’s segments age estimate decreased by ∼ 60% for segments ≤ 0.5cM compared to the current approach, Fig. 1 **E**). FastSMC’s increased accuracy in estimating coalescence times in IBD segments will translate in improved resolution for downstream applications that leverage this type of information.

### IBD sharing and population structure in the United Kingdom

We leveraged the scalability and accuracy of FastSMC to analyze 487,409 phased British samples from the UK Biobank, obtaining a fine-grained picture of the genetic structure of the United Kingdom. We detected ∼214 billion IBD segments shared within the past 1,500 years, with around 75% of all pairs of individuals sharing at least one common ancestor within the past 50 generations (Supplementary Fig. 7). Analysing the fraction of genome covered by IBD segments, we observed that 93% of individuals in the cohort have more than 90% of their genome covered by at least one IBD segment in the past 50 generations. In contrast, only ∼4% of individuals have more than 90% of their genome covered by at least one IBD segment in the past 10 generations (Supplementary Fig. 8 **A**). Looking for geographic patterns, we noticed that, despite the large sample size of the UK Biobank cohort, the average fraction of genome covered by at least one IBD segment is substantially heterogeneous across UK postcodes for recent time scales - ranging from 53.4% in London Eastern Central (EC) to 76.2% in Stockport (SK) for 10 generations - but more uniform at deeper time scales - ranging from 96.7% in London EC to 99.2% in Stockport for 50 generations (Supplementary Fig. 8 **B, C, D**). The observation of a non-uniform IBD coverage has implications for downstream methods that rely on distant relatedness at each genomic site, such as variant discovery, phasing, and imputation (see Discussion).

We analysed the network of recent genetic relatedness for 432,968 samples for whom birth coordinates are available and we constructed a 432,968 × 432,968 symmetric genetic similarity matrix where each entry corresponds to the fraction of genome shared by common ancestry in the past 10 generations for a pair of individuals. Results from agglomerative hierarchical clustering on the largest connected component of the similarity matrix (see Methods) are shown in Fig. 2. As observed in previous studies of fine-scale genetic structure (27) with a considerably smaller sample size (28), genetic clusters within the UK tend to be localized within geographic regions. Leveraging FastSMC’s accuracy and the large sample size of the UK Biobank dataset we were able to zoom into increasingly smaller regions, finding that such clusters extend beyond broad geographic clines (Fig. 2). Smaller geographic regions revealed increasingly fine-grained clusters of individuals born within a few tens of kilometers from each other, likely reflecting the presence of extended families which experienced limited migration during recent centuries. To quantify the relationship between geographic and genetic proximity, we defined 120 regions using postcodes (see Methods) and found that individuals throughout the UK, including cosmopolitan regions, find overwhelmingly more recent genetic ancestors within their own postcode than in other regions of the country, reflecting isolation-by-distance due to limited migration across the country in recent generations (see Supplementary Fig. 9 for results within the past 300 and 1,500 years, and https://ukancestrymap.github.io/ for an interactive website displaying these results). For instance, within the past 10 generations (or ∼300 years), two individuals born in North London (N) share on average 0.0092 common ancestors and two individuals born in Birmingham (B) share on average 0.0043 common ancestors. In contrast, and despite the relative geographic proximity, an individual born in North London shares on average a substantially lower 0.00059 ancestors with one born in Birmingham. We further visualized the strong link between genetic and physical distances (Supplementary Fig. 10) by building a low-dimensional planar representation of pairwise genetic distances across postcodes within the past 600 years (Supplementary Fig. 11 **B**), which we found to closely reflect geographic distance across these regions.

**Fig. 2.**
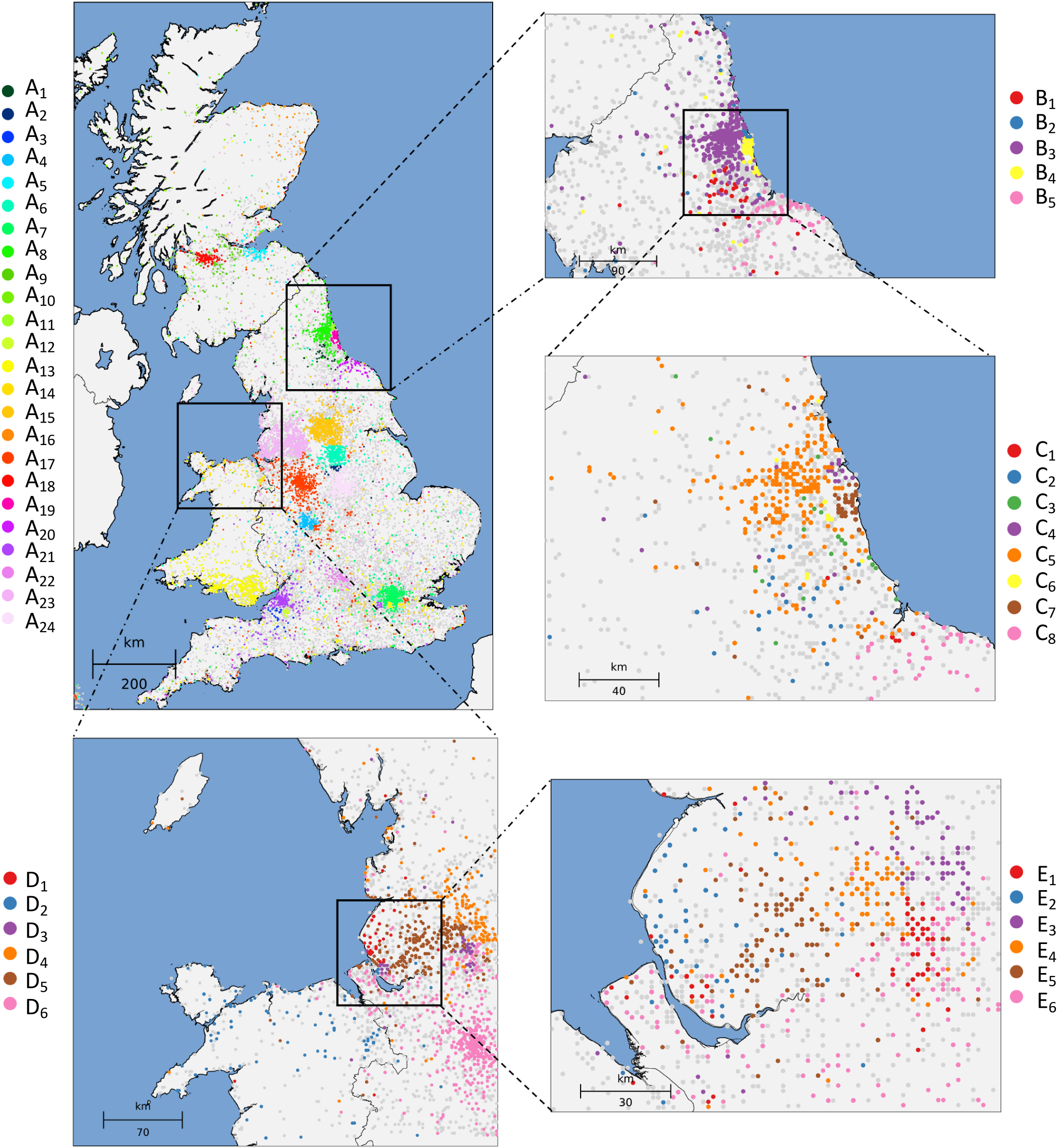
Fine-scale population structure in the UK. Hierarchical clustering of 432,866 individuals from the UK Biobank dataset based on the sharing of IBD segments within the past 10 generations. Individuals in clusters with less than 500 samples are shown in light gray. We observed 24 main clusters across the country **(top left)** and we refined two regions, corresponding to Newcastle (NE) **(top right)** and Liverpool (L) **(bottom)**, revealing fine scale population structure. No relationship between clusters is implied by the colours or cluster labelling across different plots (details are provided in Methods).

We hypothesized that the presence of such a fine-scale genetic structure in the UK may be used to effectively predict the birth location of an individual. This would imply that FastSMC may be used to predict other subtle environmental covariates, which may be causing confounding in genome-wide association studies (29). Using a simple machine learning approach (K-nearest-neighbors, see Methods and Supplementary Fig. 12), we predicted the birth coordinate of a random sample using the average birth location of the individuals we identified to be genetically closest. To increase the complexity of this task, we excluded individuals with very recent genetic ties (≤3rd degree relatives, e.g. first degree cousins (2)). We found that even if close relatives are not considered, our approach is able to predict the birth coordinates of a random sample with an average error of 95 km (95% CI=[93,97]). A standard estimate of kinship based on genome-wide allele sharing, on the other hand, achieved a considerably higher average error of 137 km (95% CI=[135,139], see Methods). Birth coordinates predicted using IBD sharing were strongly correlated to true coordinates (r=0.74, 95% CI=[0.73,0.75], for Y-coordinates and r=0.6, CI=[0.59,0.62], for X-coordinates), substantially higher than the correlation achieved using the allele sharing-based estimate of kinship (r=0.43 for Y-coordinates, 95% CI=[0.41,0.45], and r=0.31, 95% CI=[0.30,0.33], for X-coordinates). To further dissect the connection between genetics and geography in the UK Biobank, we measured the physical distance between individuals as a function of their predicted genetic and genealogical relationship. We computed the fraction of individuals who find at least one close genetic relative within the UK Biobank dataset, as shown in Fig. 3 **A**. We observed that almost all individuals (99.8%) find a genetic relative with IBD sharing equivalent to a 5th degree cousin (3.5 cM) or closer relationship, with 64.6% of samples finding a putative 3rd degree cousin (56.6 cM) or closer relative. Furthermore, stronger genetic ties translate into greater proximity of birth locations as shown in Fig. 3 **B**. For instance, for individuals sharing a fraction of genome equivalent to 3rd degree cousin or closer, the median distance between birth locations is 17 km. Very close genetic relationships are also pervasive in the dataset: about one in four individuals (23.4%) has a relative with genetic sharing equivalent to a 2nd degree cousin (226.5 cM) or closer; for these samples the median distance between birth locations is only 5 km. Additional details are shown in Supplementary Fig. 13. These findings provide empirical support to recent hypotheses that extensive segment sharing within genealogical databases may be used to recover the genotypes of target individuals (30), or to re-identify individuals through long-range familial searches (31).

**Fig. 3.**
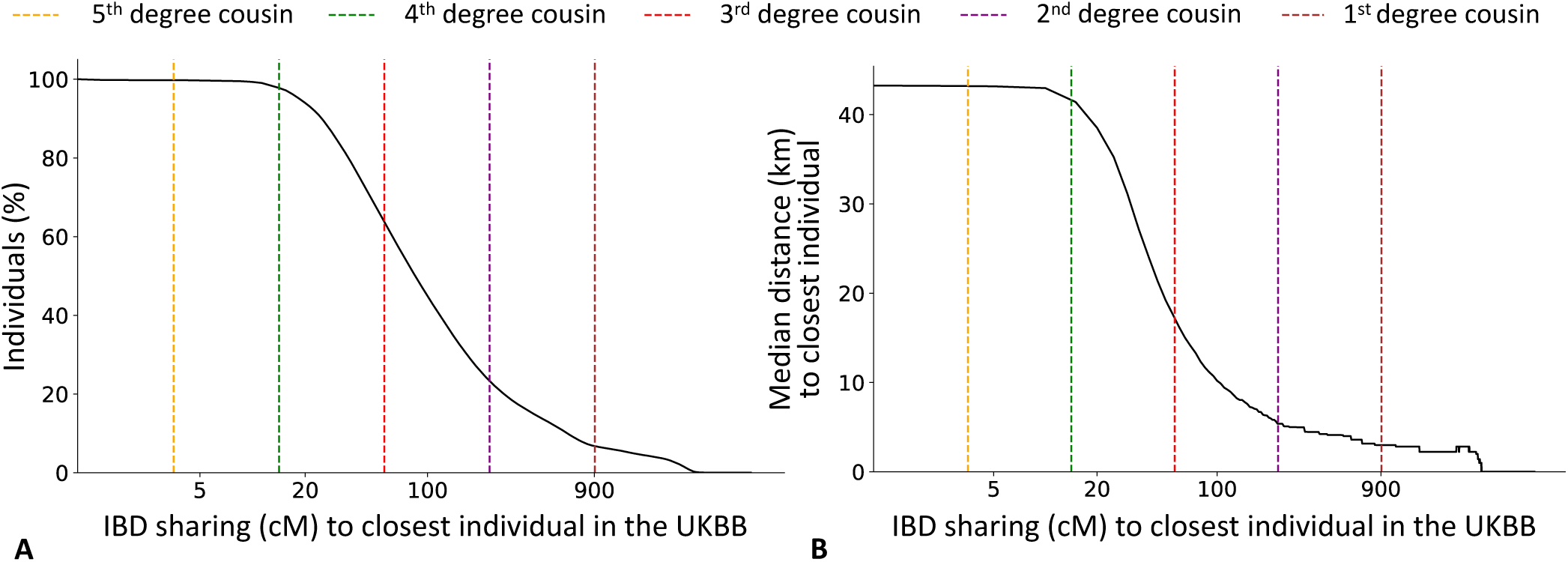
Genetic relatedness and geographic distances in the UK Biobank dataset. For each of the 432,968 UK Biobank samples with available geographic data, we detected the individual sharing the largest total amount (in cM) of genome IBD within the past 10 generations (referred to as “closest individual”). **A.** For each value *x* of total shared genome (in cM) on the X-axis, we report the percentage of UK Biobank samples (Y-axis) that share *x* or more with their closest individual. **B.** For each value *x* of total shared genome (in cM) on the X-axis, we report the median distance (km, computed every 10 cM) for all pairs of (sample, closest individual) who shared at least *x*. Vertical dashed lines indicate the expected value of the total IBD sharing for *k*-th degree cousins, computed using 2*G*(1*/*2)^2(*k*+1)^, where *G* = 7247.14 is the total diploid genome size (in cM) and *k* represents the degree of cousin relationship (e.g. *k* = 2 for second degree cousins, separated by 2(*k* + 1) generations) (10).

Analyzing broader patterns of IBD sharing, we found that individuals living in the North of the country (corresponding to Scotland and North of England) share more common ancestry than in the South, and that more generally regions within Scotland, England, and Wales tend to cluster with other regions within the same country. We estimated the effective population size from 300 years ago within each postcode (see Methods) and detected significant correlation (r = 0.28, 95% CI=[0.09,0.47] by bootstrap using postcodes as resampling unit) with present-day population density (census size per hectare), as shown in Supplementary Fig. 11 **A**. As we look deeper in time, IBD sharing patterns tend to shift and reflect historical migration events within the country. Notably, we find that individuals throughout England share deep genealogical connections with other individuals currently living in the North West and the North of Wales (Supplementary Fig. 14). These regions correspond to the “unromanised regions” of the UK and Britons living there are believed to have experienced limited admixture during the Anglo Saxon settlement of Britain occurring at the end of the Roman rule in the 5th century (32). Elevated IBD sharing between these regions may thus reflect deep genealogical connections within the ancient Briton component of modern day individuals, which is overrepresented in the North-West of the country. As expected, large cosmopolitan regions display substantially more uniform ancestry across the UK. London, in particular, has a uniform distribution of ancestry at deep time scales (50 generations, Supplementary Fig. 9), suggesting that it has attracted substantial migration for extended periods of time.

### Signals of recent positive selection in the UK Biobank

We analyzed locus-specific patterns of recent shared ancestry, seeking evidence for recent positive selection by identifying loci with unusually high density of recent coalescence times in the UK Biobank dataset. We computed the DRC_50_ (Density of Recent Coalescence) statistic (22), capturing the density of recent coalescence events along the genome within the past 50 generations, averaged within 0.05 cM windows (see Methods and Supplementary Fig. 15). Large values of the DRC_50_ statistic are found at loci where a large number of individuals descend from a small number of recent common ancestors, a pattern that is likely to reflect the rapid increase in frequency of a beneficial allele due to recent positive selection. Although, as expected, the DRC_50_ statistic computed in this analysis is strongly correlated (*r* = 0.67) with the DRC_150_ statistic that was computed using fewer samples from a previous UK Biobank data release in (22), the DRC_50_ statistic reflects more recent coalescence events than the DRC_150_ statistic, and thus more specifically reflects natural selection occurring within recent centuries.

Analyzing the distribution of the 52,003 windows in the UK Biobank dataset, we detected 12 genome-wide significant loci (at an approximate DRC_50_ p < 0.05*/*52,003 = 9.6 × 10^−7^; Fig. 4 and Supplementary Table 4). 5 of these loci are known to be under recent positive selection, harboring genes involved in immune response (*NBPF1* (33), *HLA* (34)), nutrition (*LCT* (35), *LDLR* (36)) and mucus production (*MUC2* (22)). We also detected 7 novel loci, harboring genes related to immune response (*MRC1*, playing a role in both the innate and adaptive immune systems (37), and *BCAM*, encoding the Lutheran antigen system, also associated with low density lipoprotein cholesterol measurement (38)), mucus production (*CAPN8*, involved in gastric mucosal defense (39)), tumor growth (*CHD1L*, associated with tumor progression and chemotherapy resistance in human hepatocellular carcinoma (40), and *BANP*, encoding a tumor suppressor and cell cycle regulator protein (41)), as well as genetic disorders (*HYDIN*, causing primary ciliary dyskinesia (42), and *EFTUD2*, causing mandibulofacial dysostosis with microcephaly (43)). We checked that these regions are not extreme in recombination rate or marker density (Supplementary Table 5).

**Fig. 4.**
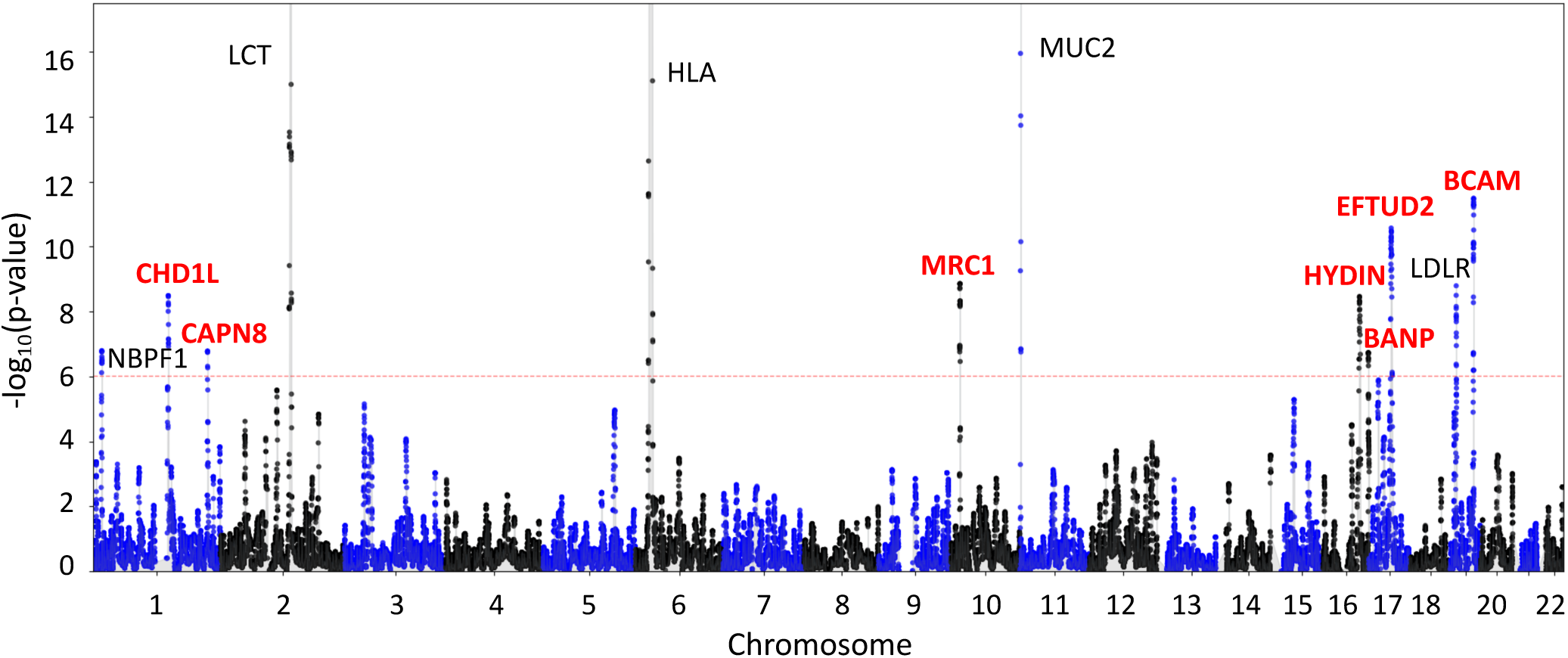
Genome-wide scan for recent positive selection in the UK Biobank dataset. Manhattan plot with candidate gene labels for 12 loci detected at genome-wide significance (adjusting for multiple testing, DRC_50_ approximate p < 0.05*/*52,003 = 9.6 × 10^−7^; dashed red line). The DRC_50_ statistic of shared recent ancestry within the past 50 generations was computed using 487,409 samples within the UK Biobank cohort. FastSMC detected 5 loci known to be under recent positive natural selection (gene labels in black) and 7 novel loci (in red).

### IBD sharing reflects sharing of ultra-rare trait-associated variation

Individuals who co-inherit a genomic region IBD from a recent common ancestor are also expected to have identical genomic sequences within that region, with the exception of de-novo mutations and other variants introduced by e.g. non-crossover gene conversion events in the generations leading to their recent common ancestor (44). We thus expect the sharing of IBD segments to be strongly correlated to the sharing of ultra-rare genomic variants (MAF< 0.0001), which tend to be very recent in origin and are usually co-inherited through recent ancestors who carried such variants. We verified this by testing for correlation between the sharing of IBD segments at various time scales and the sharing of rare variants for the ∼50*k* individuals included in the UK Biobank’s initial exome sequencing data release (45) (Supplementary Fig. 16 **A**). Specifically, we analyzed mutations that are carried by *N* out of 99,920 exome-sequenced haploid genomes (for 2 < *N* < 500), which we refer to as *F*_*N*_ mutations (46, 47). We compared the sharing of *F*_*N*_ mutations to the sharing of IBD segments in the past 10 generations within all postcodes (Fig. 5 **A**). We found that there is indeed a strong correlation between the per-postcode sharing of ultra-rare variants and the per-postcode sharing of ancestors within the past 10 generations (e.g. r = 0.3, 95% CI=[0.22,0.37] for *F*_3_ mutations found in 3 out of 99,920 chromosomes). The correlation between IBD sharing and *F*_*N*_ variant sharing decreases as *N* increases, with slightly higher correlation for more recent IBD segments, while deeper IBD sharing (within 50 generations) tends to provide better tagging of ultra-rare variants of slightly higher frequency (e.g. bootstrap p < 0.05*/*50 = 0.001 for *N* = 20; Supplementary Fig. 16 **B**).

**Fig. 5.**
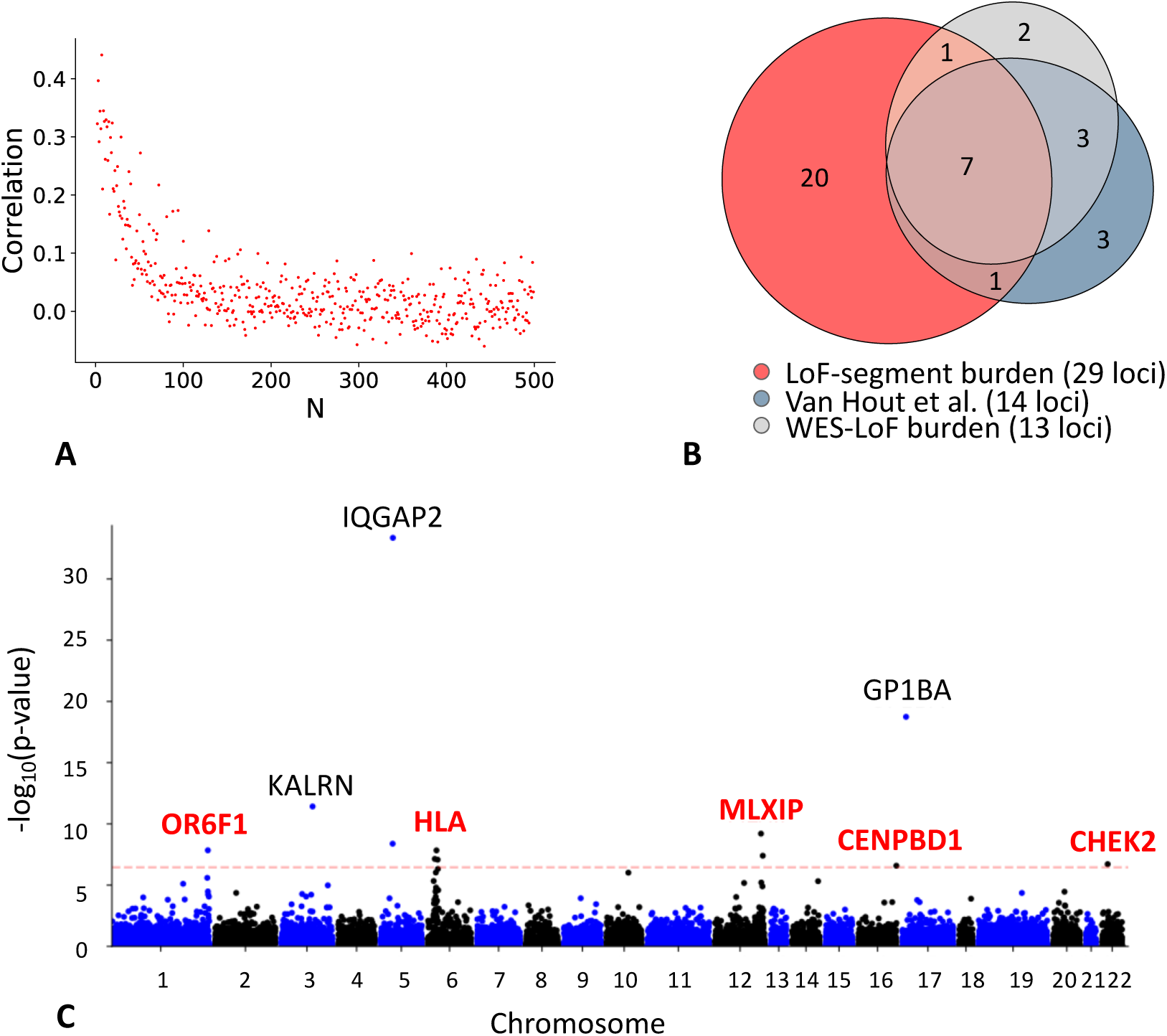
IBD sharing and rare variant associations. **A.** Correlation between IBD sharing (average number of IBD segments per pair across UK postcodes in the past 10 generations in the UK Biobank’s 487,409 samples) and ultra-rare variants sharing (average number of *F*_*N*_ mutations per pair across UK postcodes in the UK Biobank 50k Exome Sequencing Data Release for increasing value of *N*). **B.** Venn diagram representing the sets of exome-wide significant associated loci for 7 blood-related traits using three methods: the WES-based loss-of-function burden test reported by (45) (Van Hout et al.), a WES-based loss-of-function burden test we performed (WES-LoF burden), and the IBD-based loss-of-function burden test we performed (LoF-segment burden). **C.** Exome-wide Manhattan plot for mean platelet (thrombocyte) volume, after SNP-correction, using 303,125 unrelated UK Biobank samples not included in the exome sequencing cohort. Labelled genes are exome-wide significant after adjusting for multiple testing: t-test p < 0.05*/*(14,249 × 10) = 3.51 × 10^−7^; dashed red line. Black labels indicate genes that were previously reported in (45) (*KALRN, GP1BA* and *IQGAP2*), while red labels indicate novel associations detected by our LoF-segment burden analysis.

Based on this correlation between sharing of mutations and sharing of IBD segments, we hypothesized that IBD sharing of disease causing mutations would be predictive of disease. In particular, individuals who share an IBD segment with a pathogenic rare variant carrier in a known gene have a higher probability of carrying the pathogenic variant (by inheriting it from the shared ancestor) than the general population, and would thus be at increased risk for the phenotypic effect. The UK Biobank exome pilot (45) identified multiple rare coding variant burden associations with complex phenotypes, some of which were recently replicated (48, 49). We set out to test whether our IBD inference could empower us to replicate and refine these associations using the larger non-sequenced cohort. For each previously reported gene-phenotype association, we identified all sequenced individuals with a rare loss-of-function (LoF) variant (mirroring the definition of LoF in (45), see Methods) and any IBD segments they shared with individuals in the non-sequenced cohort (we refer to these as putative “LoF-segments”, though they will also include sharing of the non-LoF haplotypes because the phase of the LoF is unknown). Note that the majority of these variants are singletons or doubletons (45) and would be excluded from imputation by most current algorithms (50). Then, in the (independent) non-sequenced cohort, we tested for association between carrying a LoF-segment (the LoF-segment burden) and the phenotype known to be associated with that gene (see Methods). This approach would be optimal when IBD individuals carry a LoF variant that arose prior to the TMRCA of their shared segment. We thus tested ten transformations of the LoF-segment metric for association with phenotype to model uncertainty about the age distribution of the underlying causal variants (see Methods).

Using our LoF-segment burden, we replicated 11 out of 14 previously reported (45) associations with 7 blood-related traits at *p* < 0.05*/*10 = 0.005 (adjusted for testing of 10 transformations; see Table 1). Strikingly, we found 8 of these associations to be exome-wide significant in the non-sequenced cohort (t-test p < 0.05*/*(10 × 14,249), reflecting 14,249 genes tested using 10 transformations, Fig. 5 **B**). We next aimed at quantifying how effective IBD sharing (through LoF-segment burden testing) is at detecting associations, compared to testing directly based on exome sequencing data. We computed the phenotypic variance explained by the indirect IBD-based test and the direct exome-based test (after subtracting the effect of covariates from both, see Methods), focusing on the 14 loci reported in (45). The ratio of these variances was 19.64% on average, corresponding to the decrease in effect-size (in units of variance) due to estimation error and inclusion of segments sharing the non-LoF haplotype. We note that, due to phase uncertainty, we expect LoF-segment burden to explain at most 50% of the variance explained by direct sequencing. Assuming the ratio of variances corresponds to the squared correlation between the LoF-segment burden estimate and the true exome burden, the LoF-segment burden estimator has statistical power equivalent to a direct exome study of 19.64% of the 303,125, or ∼60*k* genotyped samples (54) - effectively doubling the size of the exome study. This demonstrates FastSMC’s accuracy and, more broadly, the utility of leveraging distant relatedness in identifying disease associations.

**Table 1.** Comparison between association analyses. We report association statistics for 14 loci and 7 traits as detected by Van Hout et al. (45) (obtained using a linear mixed model), our whole-exome sequencing burden analysis (two-sided t-test; labelled as WES LoF burden); and the LoF-segment burden (two-sided t-test). The Bonferroni-corrected exome-wide significance threshold for the first two approaches is 3.4 × 10^−6^, after correcting for multiple testing with ∼15k genes, and 3.51 × 10^−7^ for the LoF-segment burden, after adjusting for 14,249 genes and 10 time transformations. We identify 10 genes at exome-wide significance with the WES-LoF burden test, and we replicate 11*/*14 at *p* < 0.05*/*10 = 0.005 using the LoF-segment association in non-sequenced samples (8 at exome-wide significance). The last column estimates the proportion of the phenotypic variation (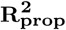, in %) of the sequenced samples that can be explained by the non-sequenced cohort (see Methods); on average that is 19.64% for all the 14 reported associations, or 27.35% if focusing on the exome-wide significant signals.

| | Gene | Trait | Van Hout et al. | WES LoF burden | LoF-segment burden | $R^2_{\text{prop}}$ |
| --- | --- | --- | --- | --- | --- | --- |
| 1 | *IL33* | Eosinophil count | 3.30E-10 | 2.01E-03 | 8.64E-15 | 72.26 |
| 2 | *GP1BA* | Mean platelet (thrombocyte) volume | 6.40E-08 | 8.84E-08 | 1.82E-19 | 32.57 |
| 3 | *TUBB1* | Platelet distribution width | 2.50E-23 | 7.34E-18 | 7.38E-12 | 07.25 |
| 4 | *TUBB1* | Mean platelet (thrombocyte) volume | 2.40E-08 | 3.01E-07 | 2.15E-03 | 04.11 |
| 5 | *TUBB1* | Platelet count | 2.10E-09 | 7.45E-07 | 4.21E-05 | 07.84 |
| 6 | *HBB* | Red blood cell distribution width | 5.80E-08 | 3.49E-02 | 2.25E-03 | 23.99 |
| 7 | *HBB* | Red blood cell count | 1.70E-09 | 7.95E-02 | 2.68E-02 | 18.23 |
| 8 | *KLF1* | Red blood cell distribution width | 1.50E-13 | 6.95E-13 | 3.49E-34 | 32.99 |
| 9 | *KLF1* | Mean corpuscular haemoglobin | 1.70E-16 | 9.11E-15 | 6.79E-21 | 16.76 |
| 10 | *ASXL1* | Platelet distribution width | 4.70E-09 | 1.44E-06 | 0.16E-00 | 00.98 |
| 11 | *ASXL1* | Red blood cell distribution width | 2.40E-11 | 8.23E-04 | 0.32E-00 | 01.03 |
| 12 | *KALRN* | Mean platelet (thrombocyte) volume | 2.70E-23 | 3.85E-18 | 3.79E-12 | 07.33 |
| 13 | *IQGAP2* | Mean platelet (thrombocyte) volume | 1.10E-19 | 3.72E-15 | 4.40E-34 | 27.43 |
| 14 | *GMPR* | Mean corpuscular haemoglobin | 1.10E-08 | 2.94E-06 | 7.60E-11 | 22.18 |

Motivated by these results, we next expanded our study to all sequenced genes for this same set of primarily blood-related traits. We identified a total of 186 exome-wide significant gene associations (t-test p < 0.05*/*(10 × 14,249)) spanning 33 genomic loci in the non-sequenced cohort by only leveraging LoF-segments; these genes were not significant in the exome burden analysis due to insufficient statistical power (see Table 1 for a comparison). We noticed that some loci included multiple significant associations, suggestive of correlation between associated features as is often seen for GWAS signals. We hypothesized that, in some cases, the true underlying causal variant may be better tagged by known high-frequency SNPs, which are likely to have been detected in previous GWAS analyses. Repeating this analysis including previously associated common variants as covariates reduced the number of significant associations to 111 across 29 loci, suggesting that inclusion of significant common associations in rare variant burden tests may lead to improved interpretability and fewer false-positives due to tagging. Results from this analysis are shown in Fig. 5 **C** and summarized in Table 2 (additional details can be found in Supplementary Fig. 17, Supplementary Fig. 18, Supplementary Fig. 19, Supplementary Fig. 20, Supplementary Fig. 21, and Supplementary Fig. 22). Our exome-wide significant signals include the association between platelet count and *MPL* (*p* = 1.99 × 10^−7^), which encodes the thrombopoietin receptor that acts as a primary regulator of megakaryopoiesis and platelet production and has not been previously implicated by genome-wide scans of either rare or common variants. We also detect several associations in genes that were not detected using exome sequencing but have been previously implicated in genome-wide scans for common variants, including associations between eosinophil count, *GFI1B* (*p* = 1.92 × 10^−7^) and *RPH3A* (*p* = 7.63 × 10^−14^) (51, 52), and associations between platelet count, *IQGAP2* (*p* = 6.52 × 10^−8^) and *GP1BA* (*p* = 1.43 × 10^−7^) (51, 53). We also identify genes that were previously associated with other blood-related phenotypes in other populations, including the association between platelet distribution width and *APOA5* (*p* = 1.94 × 10^−8^), a gene that encodes proteins regulating the plasma triglyceride levels; common variants linked to this gene have been linked to platelet count in individuals of Japanese descent (55). We detect association between red blood cell distribution width and *APOC3* (*p* = 3.67 × 10^−11^), a gene encoding a protein that interacts with proteins encoded by other genes (*APOA1, APOA4*) associated with the same trait. The association between *APOC3* and platelet count was also detected with our WES-LoF burden analysis (*p* = 2.13 × 10^−7^) and by previous studies based on common SNPs (51). We also found an association between *CHEK2* and both mean corpuscular haemoglobin (*p* = 1.43 × 10^−7^) and mean platelet volume (*p* = 1.93 × 10^−7^). This gene plays an important role in tumor suppression and was found to be associated with other blood traits, such as platelet crit, using both exome sequencing (45) and common GWAS SNPs (51), and red blood cell distribution width (52). Overall, this analysis highlights the utility of applying FastSMC on a hybrid sequenced/genotyped cohort to identify novel, rare variant associations and/or characterize known signals in larger cohorts.

**Table 2.**
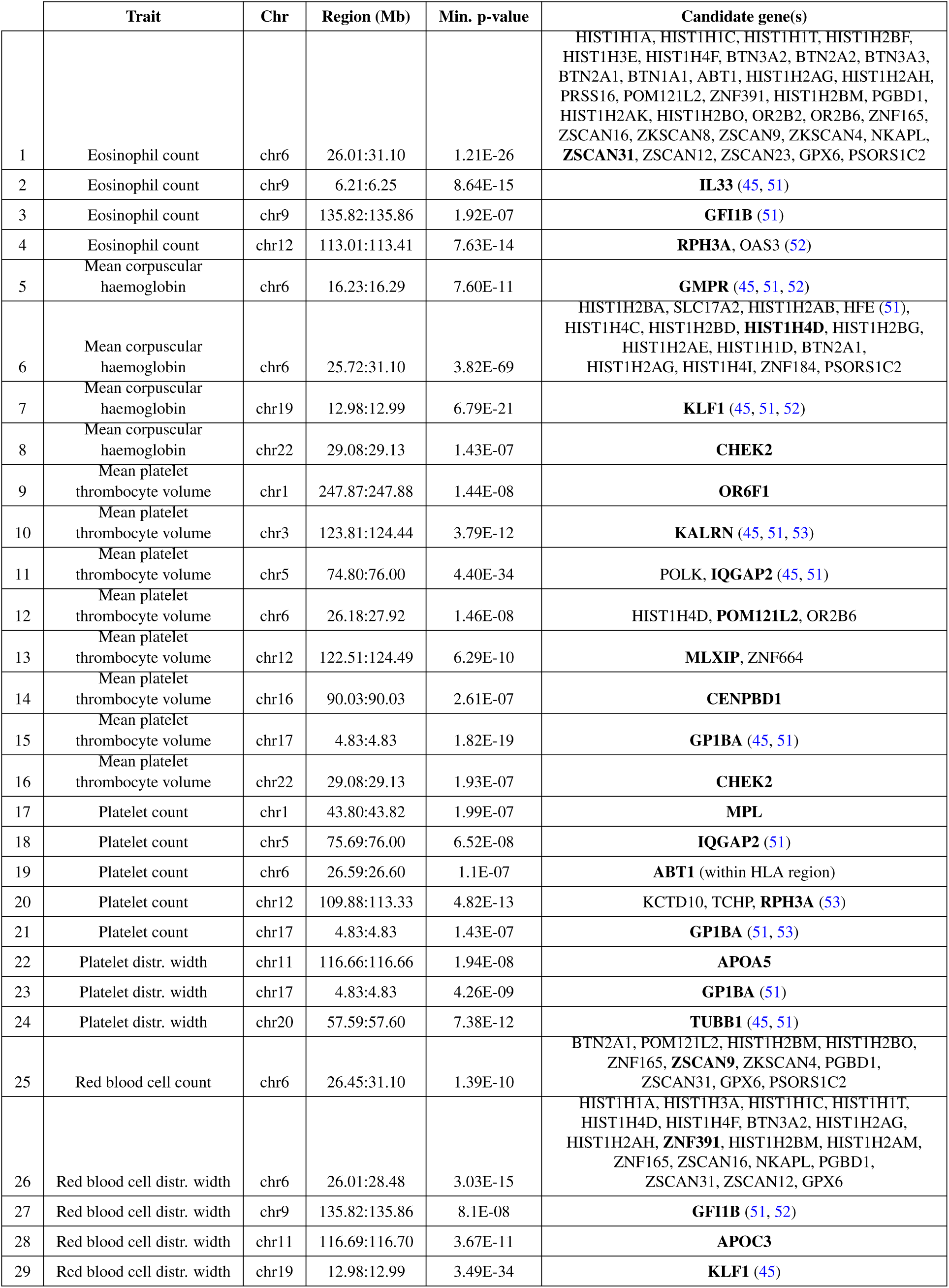
Associations detected using LoF-segment burden. Exome-wide significant associations (p < 0.05*/*(14,249 × 10) = 3.51 × 10^−7^) detected using LoF-segment burden (SNP-adjusted). Associated genes are clustered in 29 loci. For each locus we report the set of associated genes and minimum p-value. The gene corresponding to the minimum p-value is highlighted in bold.

## Discussion

We developed FastSMC, a new algorithm for IBD detection that scales well in analyses of very large biobank datasets, is more accurate than existing methods, and enables estimating the time to most recent common ancestor for IBD individuals. We leveraged FastSMC to analyze 487,409 British samples from the UK Biobank dataset, detecting ∼ 214 billion IBD segments transmitted by shared common ancestors in the past ∼ 1,500 years. This enabled us to obtain high-resolution insight into recent population structure and natural selection in British genomes. Lastly, we used IBD sharing between exome sequenced and non-sequenced samples to infer the presence of loss-of-function variants, successfully replicating known burden associations with 7 blood-related complex traits and revealing novel gene-based associations.

The level of geographic granularity that could be captured by the IBD networks emerging from our analysis underscores the importance of modeling distant relationships in genetic studies. Indeed, detecting IBD segments among close and distant relatives is a key step in many analyses, such as genotype imputation or haplotype phase inference (4, 13, 14), haplotype-based associations (11, 12), as well as in the estimation of evolutionary parameters such as recombination rates (56, 57), or mutation and gene conversion rates (44, 58). More broadly, the observed geographic heterogeneity in IBD sharing and fine-scale structure is a reminder that human populations substantially deviate from random mating, even within small geographic regions. In particular, efforts to generate optimal sequenced reference panels for imputation (59) may be greatly improved by directly sampling based on distant relatedness. Our findings also provide empirical support to recent hypotheses that relatedness within available genomic databases is sufficiently pervasive to enable recovering the genotypes of target individuals (30), or to re-identify individuals through familial searches (31). As demonstrated by our analysis, leveraging IBD to impute ultra-rare variant burden from small sequenced reference panels can be an effective approach to detect novel gene associations in complex traits and diseases. Although our analysis only considered association to individual genes, in principle genome-wide IBD sharing could also be leveraged to enhance common SNP-based risk prediction, which is particularly relevant for non-European cohorts where sequencing is limited.

FastSMC inherits some of the limitations and caveats of the GERMLINE and ASMC algorithms. First, as with most IBD detection methods, FastSMC requires the availability of phased data. We note, however, that when very large cohorts are analyzed, long-range computational phasing results in high-quality haplotype estimation with switch errors rates as low as one every several tens of centimorgans (4, 13). This is particularly true in regions that harbor IBD segments, which are of interest in our analysis. Nevertheless, our analysis has shown the presence of substantial regional heterogeneity in the extent to which individual genomes are spanned by IBD segments, which is likely reflected in a heterogeneous quality of phasing, as well as other downstream analyses, such as imputation, that rely on the presence of IBD segments. Second, FastSMC requires the input of a demographic model and allele frequencies as a prior in order to accurately estimate the age of IBD segments, and is thus subject to biases whenever the demographic model is misspecified. Although we have verified that these biases are not substantial, future work may enable us to simultaneously estimate IBD sharing and demographic history. Like ASMC, FastSMC may tolerate reasonable levels of model misspecification, but a user should be aware that issues such as substantial inaccuracies in the genetic map or strong heterogeneity of genotyping density or quality may lead to biases. Third, FastSMC currently does not enable analysis of imputed data, a limitation that is shared by other IBD detection methods, and we plan to enable this kind of analysis in future extensions of the algorithm. Finally, the accurate identification of extremely short IBD segments (<1cM) spanning hundreds of generations remains a challenge, both computationally (as the number of such segments increases very rapidly) and methodologically (as fewer variants are available to provide signal for distinguishing IBS from IBD).

In addition to algorithmic improvements to address the limitations above, we believe there are a number of interesting future extensions and interactions with other existing methods in this area. FastSMC’s segment identification step currently relies on the GERMLINE2 genotype hashing strategy. It will be interesting to test other heuristic strategies for rapidly identifying identical segments, such as the locality-sensitive hashing strategy recently implemented in the iLASH algorithm and described in a preprint as an efficient alternative to GERMLINE for segment identification (exhibiting 95% concordance with GERMLINE in application to real multi-ethnic data (60)), or methods that rely on the positional Burrows-Wheeler transform (PBWT) data structure (16, 61, 62). Several methods now exist to reconstruct gene genealogies in large samples (63–66). Two recent methods substantially improved the scalability of this type of analysis, but they either focus on data compression, relying on fast heuristics to achieve scalability at the cost of deteriorating accuracy in sparse array data (65), or employ further modeling that requires sequencing data (66), with a computational cost that is quadratic in sample size (the same computational complexity required to run the full ASMC algorithm on all sample pairs). It will be interesting to explore possible synergies between these approaches and FastSMC in large-scale genealogical analysis. Although in this work we focused on large modern biobanks comprising SNP array data, sequencing datasets are quickly becoming available. FastSMC may be tuned to enable the analysis of sequencing data as well. Finally, looking at downstream applications, a direction of future work will be to leverage FastSMC to better control for subtle population stratification for both rare and common variants in association studies. Our results show that birth coordinates can be effectively inferred from recent IBD sharing, and suggest that this may be a path towards capturing subtle environmental covariates (29) that are missed by genome-wide IBS-based approaches. FastSMC’s output could thus account for subtle stratification even when the non-genetic confounder has a small and sharp distribution, where methods such as genomic control, PCA, or mixed models have limited efficacy (67).

## Acknowledgements

We thank Po-Ru Loh, Augustine Kong, Robert Davies, Simon Myers, Geoff Nicholls, Kevin Sharp, and Brian Zhang for helpful discussions on various aspects of this work; John Watts and David Russell for discussions on historical interpretation of our results; and Fergus Cooper for help with the FastSMC software. This work was conducted using the UK Biobank resource (Application #43206). We thank the participants of the UK Biobank project. This work was supported by the MRC grant MR/S502509/1 and the Balliol Jowett Scholarship (to J.N.S.); EPSRC and MRC grant EP/L016044/1 (to G.K.); NIH grant R21-HG010748-01 (to P.F.P. and A.G.); Wellcome Trust ISSF grant 204826/Z/16/Z (to P.F.P. and M.R.); and by ERC Starting Grant 850869 (to P.F.P.).

## Methods

### FastSMC identification step

FastSMC’s identification step leverages genotype hashing, a strategy that was introduced by the GERMLINE algorithm (19) to obtain substantial gains in computational scalability in the detection of pairwise shared IBD segments. This approach restricts the search space of IBD pairs to those that have small, identical shared segments, which are then extended to long segments with some tolerance. This in turn reduces the IBD search from all pairs of individuals to the subset of pairs that produce a hash collision at a given segment plus the cost of hashing the genotype data, which is linear in sample size and genome length, thus dramatically reducing the cost of IBD detection. However, a limitation of this strategy is that certain short haplotypes can be extremely common in the population and result in hash collisions across a large fraction of samples, effectively reverting back to a nearly all-pairs analysis and monopolizing computation time (Supplementary Fig. 1, gray). These common haplotypes are likely due to recombination cold-spots and typically contain little variation to classify shared segments. The majority of computation is thus spent processing regions with the least information content. The GERMLINE2 algorithm, which we developed in this work, proposes a novel adaptive hashing approach that adjusts to local haplotype complexity to dramatically reduce computational and memory requirements. GERMLINE2 proceeds as follows: (1) the input haplotype data is divided into small windows containing 16 or 32 SNPs each (depending on memory architecture); for a given window *w*, (2) all haplotypes are converted to binary sequences and efficiently hashed into bins of identical segments; (3) for each bin that contains more individuals than a fixed threshold (i.e. a low complexity bin) step 2 is recursively performed for window *w+1* until no more low complexity bins are found; (4) all pairs of individuals sharing within a bin are then recorded in a separate hash table that stores putative segments; (5) pairs of individuals sharing contiguous windows that are sufficiently long are reported for validation. The primary computation speed-up comes from the recursive hashing step, which requires haplotypes to be sufficiently diverse before they are explored for pairwise analysis and stored (Supplementary Fig. 1, green). To allow for possible phasing errors, a putative shared segment is maintained through a parameterized number of non-identical windows, and the total number of non-identical windows within the segment is also reported for filtering. Most phasing errors either appear as ‘‘blips”, where a phase switch is immediately followed by a switch back, or by single switches followed by long stretches of accurate phase (13, 14) - both of which are permitted by allowing periodic non-matching along the putative IBD segment. This permissive treatment of phasing is further filtered in the validation step (below). GERMLINE2 thus does not require any backtracking and only a small number of physical windows need to be stored in memory at any time (only enough to perform the recursion), allowing the method to run on input data of unlimited length.

### FastSMC validation step

Every segment detected by GERMLINE2 in the identification step is added to a buffer of candidate segments. These segments are immediately decoded by the *ascertained sequentially Markovian coalescent* (ASMC) algorithm (22) once the buffer is full. ASMC is a coalescent-based HMM (23, 24, 26, 68) that estimates the posterior of the coalescence time, or TMRCA, for a pair of individuals at each site along the genome using either sequencing or SNP array platforms. It leverages a demographic model as prior on the TMRCA, which increases accuracy in detecting regions of low TMRCA (25), but would be infeasible to apply to the analysis of all pairs of genomes, and is thus only applied with the goal of validating previously identified candidate IBD regions. The hidden states of the HMM are discretized intervals, corresponding to a user-specified set of TMRCA intervals. The HMM emissions probabilities correspond to the probabilities of observing both the genotypes of the pair of analyzed individuals and the frequencies of mutations along the sequence, given the pair’s TMRCA at each site, and the frequency of the allele (26). The HMM transitions between hidden states correspond to changes in TMRCA along the genome due to recombination events, based on the conditional Simonsen–Churchill model (69, 70). The demographic history of the analyzed haplotypes is first estimated using other methods (e.g. (24, 26)) and provided in input, so that the initial state distribution, the transition, and the emission probabilities can be computed. The most likely posterior sequence of TMRCAs along the genome is inferred using a dynamic programming approach that requires computing time linear in the number of hidden states, leading to a substantial speed-up over an HMM’s standard forward-backward algorithm, which scales quadratically in the number of hidden states. When decoding the buffer of candidate segments, ASMC computes the posterior of the coalescence time for each candidate segment and each site from the minimum starting position to the maximum ending position in the buffer. At each site, if the posterior of coalescence time being between present time and the user-specified time threshold is higher than its prior, the site is considered to be IBD and the IBD segment is extended to the next site if the same condition is still satisfied, obtaining multiple IBD segments, all shorter than the original IBD candidate segments. The average probability of the TMRCA being between present time and the user-specified time threshold is computed over all sites until the segment breaks. This average probability corresponds to an IBD quality score: the higher it is, the more likely the segment is IBD. Each segment is also associated with an age estimate corresponding to the average maximum-a-posteriori (MAP) along the segment. FastSMC finally outputs each IBD segment with its corresponding IBD quality score and age estimate.

### Simulations

Unless otherwise specified, all simulations use the setup of (22), which is described in this section. We used the ARGON simulator (v.0.1.160615) (71), incorporating recombination rates from a human chromosome 2 (see URLs) and a recent demographic model for European individuals (Northern European (CEU) population (26)). For each dataset, we simulated 300 haploid individuals and a region of 30 Mb. To simulate SNP array data, we subsampled polymorphic variants to match the genotype density and allele frequency spectrum observed in the UK Biobank dataset. We used recombination rates from the first 30 Mb of chromosome 2 (average rate of 1.66 cM per Mb). No genotyping or phasing error was introduced in our simulations. We simulated one dataset following this setup to fine-tune parameters, and 10 other datasets (all with different seeds) for accuracy benchmarking. The demographic model and genetic map used to simulate the data were used when running FastSMC, unless otherwise specified. When testing FastSMC’s robustness to demographic model misspecification, we simulated data under a constant population size of 10,000 diploid individuals, but ran FastSMC assuming a European demographic model.

### Accuracy evaluation

We compared FastSMC to the most recent published software version available for existing methods at the time we conducted this analysis: germline-1-5-2, refined-ibd.23Apr18.249 and RaPID_v.1.2. We adopted the following definition of IBD sharing: at a given site along the genome, a pair of individuals is defined to be IBD if their TMRCA is below a user-specified time threshold. We benchmarked all methods using several such time thresholds (25, 50, 100, 150 and 200 generations), testing all polymorphic sites for all pairs of genomes in the simulated data, across 10 coalescent simulations. Accuracy was quantified using the area under the precision-recall curve (auPRC), which effectively addresses issues with class imbalance that are expected in this analysis due to the low prevalence of IBD sites compared to non-IBD sites. For a given IBD time threshold, a site inferred to be IBD by one of the methods was considered correct (true positive) if the true TRMCA at this site was indeed below the specified IBD time threshold, and incorrect (false positive) if the true TRMCA at this site was above the IBD time threshold. Similarly, a site that was not reported to be IBD by the tested method was considered correct/incorrect (true/false negative) if the true TMRCA at the site was found to be below/above the IBD time threshold. We used these definitions to compute the precision and recall values for all methods. Each method presents different parameters, which can be used to tune precision and recall, e.g. by allowing a more or less permissive detection of IBD segments. String-matching methods (GERMLINE and RaPID) report long, approximately IBS regions and do not produce calibrated estimates of segment quality. A commonly used proxy for the likelihood of a detected IBS segment being IBD is its length, with longer IBS segments being more likely to be IBD than shorter ones. In lack of an interpretable measure of accuracy, precision and recall for the output of GERMLINE and RaPID was thus tuned by using different segment length cut-offs. RefinedIBD and FastSMC, on the other hand, both provide an explicit quantification of segment quality. RefinedIBD outputs a LOD score for each segment, while FastSMC computes a segment’s IBD quality score, which is the posterior of the TMRCA being between present time and the user-specified time threshold. We thus used LOD score and IBD quality score to tune precision and recall of RefinedIBD and FastSMC, respectively. Each method presents a number of additional parameters, which we further optimized using a grid-search (see below), so that each method can be run with a set of parameters that is as close to optimal as possible. Despite the extensive tuning, not all accuracy values could be explored by all methods. Namely, some recall values cannot be achieved using realistic parameters, due to factors such as the minimum allowed LOD parameter for RefinedIBD, the time discretisation introduced in FastSMC and the minimum length parameter for all methods. We thus evaluated all algorithms by restricting the comparison to the range of recall values that could be achieved by all methods, which we refer to as common recall range. Furthermore, some of these parameters affect the speed of each algorithm. The parameters we chose for comparing methods are optimized for maximum accuracy, although we avoided parameters that would result in degeneracies (e.g. the minimum length in FastSMC’s identification step could be set to values below 0.1 cM, effectively disabling this step and reverting to a pairwise ASMC analysis, which would lead to a higher accuracy and larger recall range, at the cost of unreasonable computation).

### Fine-tuning of methods

For each method (FastSMC, RefinedIBD, GERMLINE and RaPID) and each IBD time threshold (25, 50, 100, 150 and 200 generations), we performed a grid-search over possible parameter values to optimise the accuracy on one simulated dataset and select the best set of parameters. For each method, we then explored the obtained set of parameters to make the algorithm faster while negligibly compromising the accuracy (in most cases, this resulted in a slightly smaller recall range but with a substantial gain in speed). We finally used 10 independent simulated datasets to validate the accuracy and the UK Biobank dataset to measure running time and memory usage (see Methods). Unless specified otherwise, the parameters presented in Supplementary Table 1 were used in all analysis.

### Computing confidence intervals

Unless otherwise indicated, confidence intervals (CIs) were computed by bootstrap using 39 genomic regions as resampling unit. The use of genomic regions as resampling unit, rather than individuals, ensures that approximately independent bootstrap replicates are utilized. These 39 genomic regions were obtained by dividing the genome (autosomal chromosomes only) in regions from different chromosomes or separated by centromeres.

### Estimating the age of an IBD segment

A common way to estimate the age of an IBD segment is to use its length to obtain a maximum likelihood estimator (MLE). When the time in generations *g* to the most recent common ancestor is known, the total length of a randomly chosen shared IBD segment follows an exponential distribution with rate 1*/*2*g* per Morgan (5). The likelihood function is thus given by 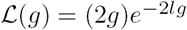, where *l* denotes the segments length in Morgans. As 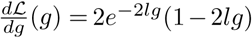, the MLE is given by 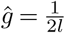. FastSMC does additional modelling and provides a different age estimate, which consists in the average maximum a posteriori (MAP) estimate of the TMRCA along the segment. We sometimes report segment age estimates in years, rather than generations, assuming 30 years per generation.

### Effective population size estimate

Assuming a constant effective size *N*_*e*_, the probability of finding a common ancestor at a given site for a pair of individuals is exponentially distributed with mean *N*_*e*_. 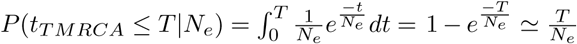 for *N*_*e*_ → ∞ i.e 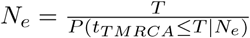 for *T* > 0. We estimate the probability of finding a common ancestor before any time threshold *T* using the fraction of genome shared by IBD segments denoted by *f*_*T*_. Let Γ denote the set of sites along the genome and *θ* the demographic model. 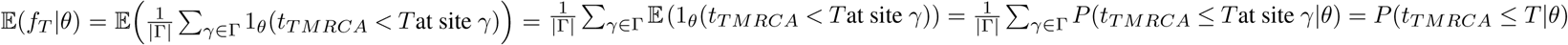. We thus estimate effective population size *N*_*e*_ using 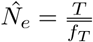 for any *T* > 0, where 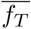 is the sample mean for the fraction of genome shared *f*_*T*_. Note that, for simplicity, we infer a single aggregate effective population size across the past *T* generations rather than comparing more complex demographic models.

### UK Biobank dataset and definition of postcodes

The UK Biobank cohort (2) contains 487,409 samples, which were phased at a total of 678,956 autosomal biallelic SNPs using Eagle2 (14). 49,960 of these individuals were also exome-sequenced resulting in ∼4 million polymorphic variants, 98.4% of which have frequency less than 1% (45). We used the same sets of unrelated individuals (*N* = 407,219) and individuals of White British ancestry (*N* = 408,974) as defined in (2). Related individuals refer here to ≤3rd degree relatives, e.g. first degree cousins, estimated using the software KING (72). FastSMC was run on the UK Biobank data set using the optimal set of parameters described earlier, using the demographic model for CEU individuals inferred in (26). We note that several candidate demographic models could be adopted (e.g. that of (8)) and that simultaneous estimation of IBD sharing and demographic model are an attractive direction of future investigation (see Discussion). We occasionally divided the genome in 39 autosomal regions from different chromosomes or separated by centromeres as previously done in (22). This enabled us to efficiently parallelize the analysis, and prevented issues due to low marker density in centromeres. Birth coordinates (all within the UK) were available for 432,968 individuals in the cohort in the Ordnance Survey Great Britain 1936 (OSGB36) Eastings and Northings system. We refer to these coordinates as X and Y coordinates, which can be converted into longitude and latitude. We analyzed population structure using 120 postcodes in the UK, only looking at the first one or two letters indicating the city or region. Postcodes BT, BF, BN and CR were not included due to lack of samples.

### Hierarchical clustering

We constructed a similarity matrix for all individuals with birth coordinates in the UK Biobank dataset (432,968 samples) using the sharing of IBD segments within the past 10 generations. This resulted in a sparse 432,968× 432,968 matrix, where entry (*i,j*) corresponds to the fraction of genome shared by common ancestry in the past 10 generations between individuals *i* and *j*. We computed the largest connected component of this matrix, which comprised all but 102 individuals, which we excluded from further analysis. We then applied an Agglomerative Hierarchical Clustering algorithm for Sparse Similarity Matrices using average linkage (sparseAHC library, see URLs). We obtained a dendrogram with 432,866 leaves (one for each sample) and a single root. Each node is annotated with a distance from the root, ranging from 0 (the root itself) to 1 (the leaves). A node’s distance from the root corresponds to the fraction of genome shared by individuals whose TMRCA is such a node. To visualize clustering of individuals in Fig. 2, we cut the tree at increasingly large distances from the root, corresponding to increasingly fine-grained clusters in terms of both genetic and geographic proximity of the samples. In each case, we only highlight large clusters containing at least 500 individuals. To plot results, we divided each submap into 10,080 grid cells (80 lines along the X-axis and 126 lines along the Y-axis). In each cell, we computed the most represented cluster (i.e the cluster with the largest number of individuals in that cell) and individuals from that cluster are shown in the corresponding color. The transparency of all points within a cell (ranging from 0 to 1) was set to the fraction of individuals from that cell corresponding to the most represented cluster. All light gray dots correspond to individuals that are either in clusters containing less than 500 samples or part of a cluster different from the one represented in the grid cell.

### Prediction of birth location

We randomly sampled two subsets of 10,000 individuals each from the UK Biobank cohort and used the sharing of recent common ancestors in the past 600 years with the remaining 412,866 individuals in the UK Biobank dataset to predict their birth locations, applying the K-nearest-neighbors algorithm. One dataset was used to find the optimal value for the parameter *K*, while the second one was used to validate the results (details shown in Supplementary Fig. 12). We computed pairwise genetic similarity across individuals using either FastSMC-estimated pairwise IBD sharing, or using a standard estimate of kinship based on genome-wide allele sharing. This kinship estimator was obtained by computing the product **XX**^⊤^, where **X** is the *N* × *S* genotype matrix (*N* = 432,866 samples and *S* = 716,175 autosomal SNPs), standardized to have mean 0 and variance 1 for each column. Sharing of very close relatives is highly informative of geographic proximity. We thus excluded ≤3rd degree relatives (2) from the dataset to bypass this source of information and test the generality of this approach. Prior to the exclusion of close relatives, the IBD-based predictor obtained an average error of 86 km (95% CI=[83,88], optimal K=1), while the allele sharing distance predictor obtained an average mean error of 118 km (95% CI=[115,121], optimal K=1). After removing close relatives (which brought sample size down to 8,226), the IBD-based predictor obtained an average error of 95 km, 95% CI=[93,97], optimal K=5 compared to 137 km, 95% CI=[135,139], optimal K=5 for allele sharing.

We regressed the true X (resp. Y) birth coordinates on the predicted X (resp. Y) birth coordinates using either IBD sharing in the past 20 generations allele sharing, for the set of 8,226 random samples we obtained after excluding 3rd degree relatives (2). The estimated coefficient for the IBD-based predictor was 0.91, 95% CI=[0.88,0.94], (resp. 0.96, 95% CI=[0.94,0.98]), substantially larger than the estimated coefficient for the allele sharing predictor (0.12, 95% CI=[0.09,0.16]) (resp. 0.12, 95% CI=[0.09,0.15]). Finally, after excluding close relatives, we computed the correlation between true and predicted coordinates for both methods, obtaining a stronger correlation when using IBD sharing (*r* = 0.6 for X coordinate and *r* = 0.74 for Y coordinate) than when using allele sharing (*r* = 0.31 for X coordinate and *r* = 0.43 for Y coordinate, respectively). Correlation for IBD sharing was also stronger without removing close relatives (*r* = 0.63 for X coordinate and *r* = 0.77 for Y coordinate, compared to *r* = 0.40 and *r* = 0.42 respectively for allele sharing).

### Detection of recent positive selection

In order to identify genomic regions with an usually high density of coalescence times, we computed the DRC_T_ (Density of Recent Coalescence) statistic within the past T generations (22). The DRC_50_ statistic reflects the probability that a random pair of individuals coalesced at a given genomic site during the past 50 generations, averaged within windows of 0.05 cM along the genome. FastSMC does not output the posterior of the TMRCA but provides an “IBD quality score”, corresponding to the sum of posterior probabilities between generations 0 and *T*, where *T* is the user-specified threshold. As the UK Biobank dataset was analysed for *T* = 50, the DRC_50_ statistic at a given site along the genome was estimated by averaging all IBD quality scores obtained from all analyzed pairs of samples (assuming a score of 0 if no segment is present for a pair). Results are presented in Fig. 4 and Supplementary Table 4. When multiple candidate genes were found, we only retained the one nearest to the top SNP (i.e with smallest p-value).

Given *n* samples from a population of recent effective size *N*, the DRC_50_ statistic is approximately Gamma-distributed under the null for *n* ≪ *N* (22). Following (22), we thus built an empirical null model using a database of regions under positive selection (see URLs). We fitted a Gamma distribution (using the Scipy library, see URLs) to the estimated DRC_50_ values within putative neutral regions (after excluding the regions of known positive selection and 500 kb windows around them), and used this model to obtain approximate p-values throughout the genome. We analyzed 52, 003 windows, using a Bonferroni significance threshold of 0.05*/*52,003 = 9.6 × 10^−7^. Three of the genome-wide significant signals we detected (*MRC1* locus, chr10:17.43-18.10 Mb; *HYDIN* locus, chr16:70.10-72.69 Mb and *EFTUD2* locus, chr17:41.84-44.95 Mb) fell within the putative neutral regions of the genome. We thus iterated this procedure, excluding these loci from the set of putative neutral loci. Once again, one of the significant genome-wide significant loci (*BANP* locus, chr16:88.25-88.48 Mb) overlapped with the putative neutral regions. We excluded this locus and iterated the procedure again. Results from the empirical null model fitting are presented in Supplementary Fig. 15. Finally, we verified that the genome-wide significant peaks detected using this approach are not found in regions of extremely high recombination rate or low marker density, which may introduce systematic biases in IBD detection Supplementary Table 5.

### Association analyses

We used each IBD segment between exome-sequenced, LoF carrying individuals and non-sequenced individuals as a surrogate for the latter carrying an untyped LoF mutation, which we then tested for association with phenotype. For a given gene and a given non-sequenced individual, we define a “LoF-segment” as any IBD segment shared with an exome-sequenced LoF mutation carrier. We then compute a LoF-segment burden for each individual as the sum of probabilities (IBD quality scores) of all LoF-segments involving that individual, under the assumption that increased IBD probability and incidence corresponds to increased probability of sharing the LoF variant. Finally, this burden is tested for association with each target phenotype (rank-based inverse normal transformed) in a linear regression with covariates for age, sex, BMI, smoking status, and four principal components, similarly to (45).

Although this test captures uncertainty about the sharing of IBD segments through the use of IBD quality scores, it makes use of all LoF-segments, regardless of their age. As a result, it may be suboptimal in cases where the LoF arose after the TMRCA, for which a LoF-segment is independent of underlying LoF sharing, and thus do not contribute signal to the burden test. We thus augmented the LoF burden test by separately considering only LoF-segments older than a specified threshold. For each gene, we divided all LoF-segments into deciles based on IBD quality score. For instance, segments with IBD quality scores in the tenth decile (which corresponds to the IBD quality score interval [0.47, 1]), strongly suggest the sharing of common ancestors that lived recently and have therefore transmitted extremely recent variation. We then constructed ten separate LoF-segment burdens, with increasingly more stringent quality score cutoffs (referred as time transformations in our analysis), and performed ten association tests for each gene, taking the test that resulted in the lowest p-value after adjusting significance thresholds by conservatively assuming independence for all tests. Not all genes contained shared LoF-segments for testing, which reduced the total number of tested genes to 14,249. This resulted in a Bonferroni-corrected exome-wide significance threshold of 0.05*/*(10 × 14,249) for our LoF-segment burden analysis.

Although gene-based burden tests are meant to implicate specific genes with a known directional effect on the trait, the observed signal may not always be driven by a causal variant, and instead be due to tagging of causal variants in nearby genes. In this case, it is possible that the underlying rare causal variant is tagged by a common variant, which may have been detected in a previous GWAS. In particular, these common variants may provide better tagging of the underlying true causal variation than our LoF-segment burden score, and would thus remove or significantly reduce the association signal if included as covariates in the test. Based on this principle, for each gene and each trait, we selected up to three genotyped SNPs that were in proximity (*±*1 Mb from the gene), which were significantly (*p* < 1 × 10^−8^) associated in (73), and used them as covariates. We observed that this approach often improves the association signal (e.g. see Supplementary Fig. 17), removing signals that were likely caused by tagging common variants. We refer to analyses that include top associated SNPs as covariates as *SNP-adjusted*, for either the LoF-segment or WES-LoF burden test; results without the SNP-adjustment are shown in Supplementary Fig. 17 and Supplementary Table 6.

We validated our approaches, both *LoF segment* and *not SNP-adjusted LoF segment*, which seek to implicitly impute ultra-rare LoF variation between sequenced and non-sequenced individuals, by testing for association between rare variation and 7 blood-related phenotypes recently analyzed in (45), and comparing to the results of that same study. However, because summary association statistics for this analysis are not available, we performed our own exome-wide burden testing. Specifically, we used the same testing framework we used in our LoF-segment burden analysis to test for association between phenotypes and burden of LoF variants within a gene in exome-sequenced individuals, adjusting for the same covariates and using the same rank-based inverse normal transformation for the phenotype. We refer to this analysis as WES-LoF burden analysis.

Both LoF-segment burden and WES LoF burden analyses were restricted to unrelated individuals of White British ancestry, as defined in (2), and the LoF-segment burden analysis was further restricted to individuals for which exome sequencing data is not available. This resulted in 303,125 individuals for the two LoF-segment burden tests and 34,422 individuals for the WES LoF. Finally, we note that the UK Biobank has recently released a statement regarding incorrectly mapped variants in the 50k WES “Functionally Equivalent” (FE) dataset (see URLs), which we however believe did not introduce any significant biases in our analyses.

Applying our WES-LoF burden analysis we detected 10 out of 14 exome-wide significant associations also reported in (45). We also detected 3 additional associations that were not reported in (45): *MAPK8* and *APOC3* with red blood cell distribution width (*p* = 1.32 × 10^−6^ and *p* = 2.13 × 10^−7^ respectively), and *TET2* with eosinophil count (*p* = 1.79 × 10^−8^). Results are summarized in Fig. 5 **B**. These differences are likely ascribed to the slightly different testing strategy we adopted, e.g. the use of a linear model, rather than a linear mixed model, and the exclusion of related samples. Detailed results for these analyses are reported in Table 1, Table 2, and Supplementary Table 6. A QQ-plot verifying the calibration of our test for the SNP-adjusted LoF-segment burden analysis is shown in Supplementary Fig. 23 and, as explained at that point, rare variant stratification is likely to be included in our results (67) but addressing this issue goes beyond the scope of the current study.

### URLs

- The FastSMC software will be made available upon publication at https://www.palamaralab.org/software/FastSMC.
- Interactive map and results from demographic analysis are available at https://ukancestrymap.github.io/.
- ARGON simulator: https://github.com/pierpal/ARGON.
- UK Biobank: http://www.ukbiobank.ac.uk/.
- Human genetic maps: http://www.shapeit.fr/files/genetic_map_b37.tar.gz.
- dbPSHP database of positive selection: ftp://jjwanglab.org/dbPSHP/curation/dbPSHP_20131001.tab
- 2011 census population size in Wales and England: https://www.nomisweb.co.uk/census/2011/qs102ew.
- Data on population in Scotland areas: https://www.scotlandscensus.gov.uk/ods-web/data-warehouse.html#bulkdatatab.
- Reference genome and gene coordinates: http://hgdownload.cse.ucsc.edu/goldenPath/hg19/database/refGene.txt.gz.
- Summary statistics from the BOLT-LMM association study: https://data.broadinstitute.org/alkesgroup/UKBB/.
- Python’s Scipy library: http://www.scipy.org/.
- Python’s sparseAHC library: https://rdrr.io/github/khabbazian/sparseAHC/.
- Note on UK Biobank exome dataset issues: https://www.ukbiobank.ac.uk/wp-content/uploads/2019/12/Description-of-the-alt-aware-issue-with-UKB-50k-WES-FE-data.pdf.

## SUPPLEMENTARY FIGURES

**Supplementary Fig. 1.**
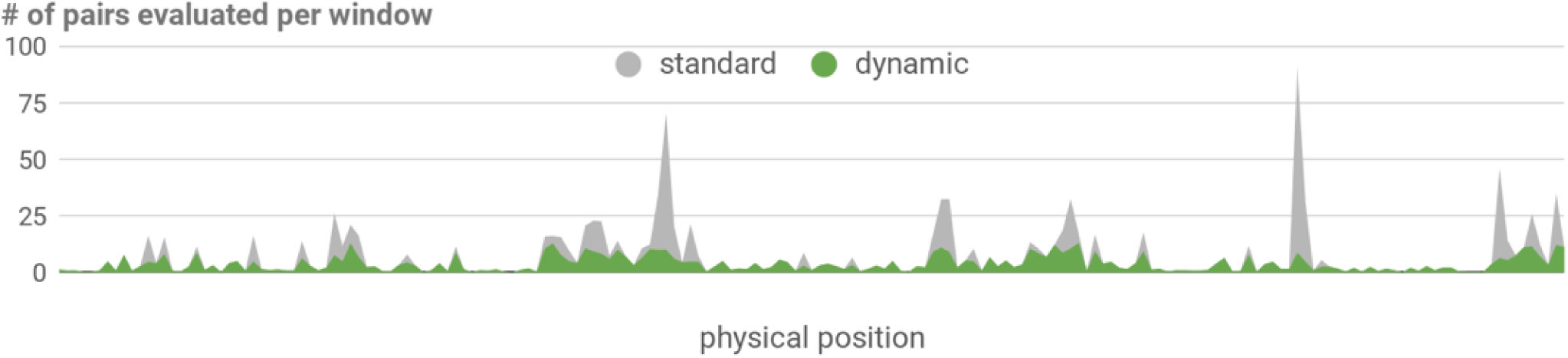
GERMLINE and GERMLINE2 comparison. Search complexity for standard vs dynamic hashing. Number of pairs of individuals (in millions) with identical haplotypes in each genome window using GERMLINE standard hash (gray) and the proposed GERMLINE2 dynamic hash (green). Analysis of 16,000 random UK Biobank samples from chromosome 22. Multiple regions of the genome exhibit sharing between >10% of all pairs, requiring extensive follow-up analysis in standard mode.

**Supplementary Fig. 2.**
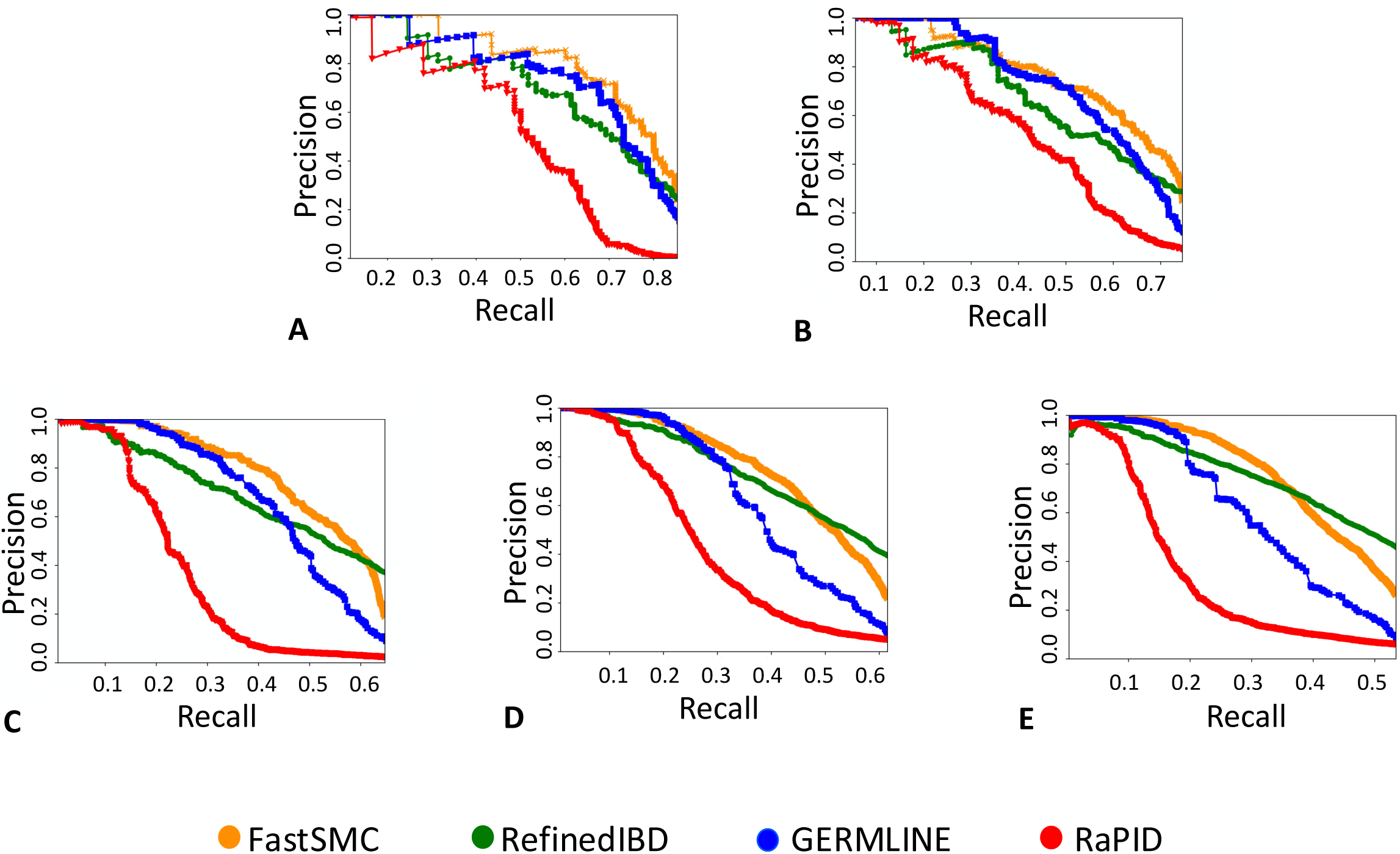
Accuracy evaluation for IBD detection at different time scales. Precision-recall curves within the common recall range (i.e where all methods are able to provide predictions) in the past 25 (**A**), 50 (**B**), 100 (**C**), 150 (**D**) and 200 (**E**) generations, randomly sampled from 10 realistic simulated datasets. Each dataset consists of a 30Mb chromosome under European demographic history model, recombination rates from a human chromosome 2 and SNP ascertainment matching UKBB allele frequencies. Precision refers to the fraction of true IBD segments among the retrieved segments while recall measures the proportion of actual IBD segments that are correctly identified as such. For each method (FastSMC, RefinedIBD, GERMLINE and RaPID), optimal parameters from the grid search were used.

**Supplementary Fig. 3.**
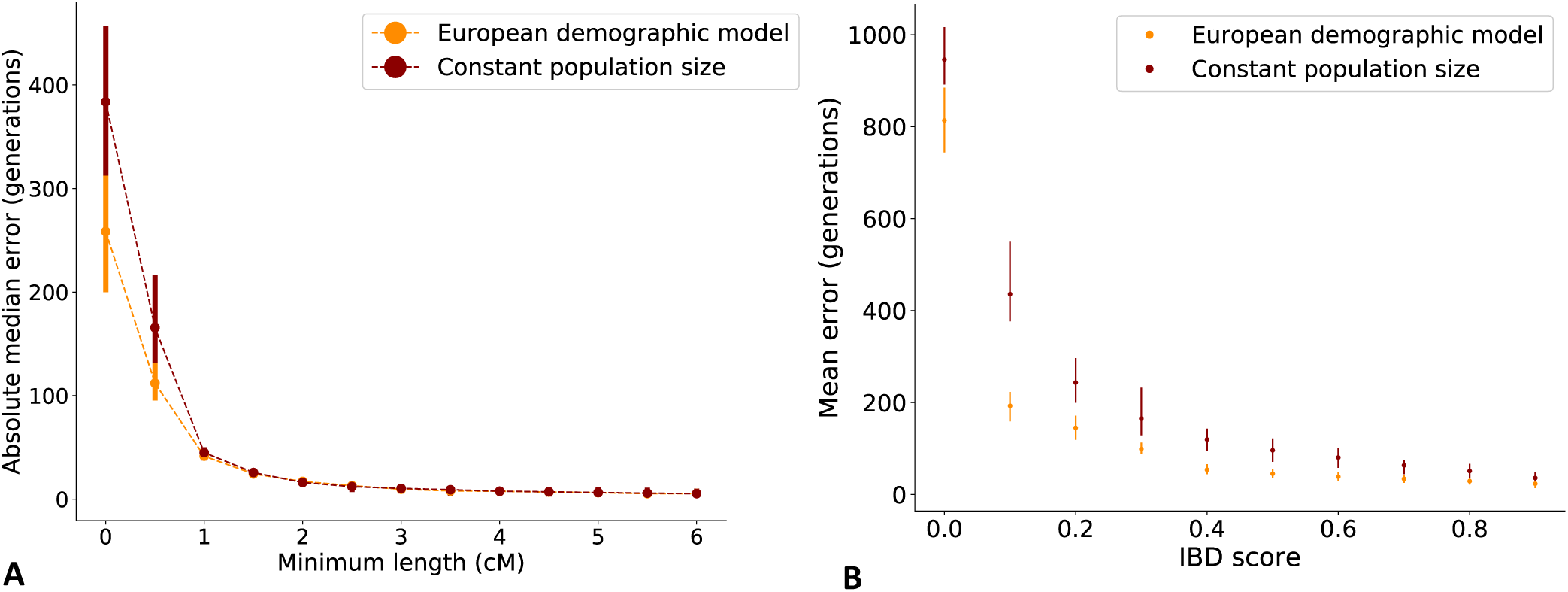
Demographic model misspecification. Effects of demographic model misspecification in age estimate of IBD segments. We simulated 10 batches of 300 haploid samples from the first 30Mb of a human chromosome 2 and a constant population size of 10,000 diploid individuals, and 10 realistic batches according to the setup described in Methods. We ran FastSMC on both datasets, assuming a European demographic model, with a time threshold of 50 generations and a minimum length of 0.001 cM. We report **(A)** the absolute median error between the true age and the MAP age estimate while varying the minimum length of the IBD segments, and **(B)** the mean error between the true age and the MAP estimate while varying the minimum IBD score. Vertical lines represent the standard error over all 10 batches. Misspecifying the demographic model results in biased age estimates for very short (< 1cM) or intermediate IBD score segments.

**Supplementary Fig. 4.**
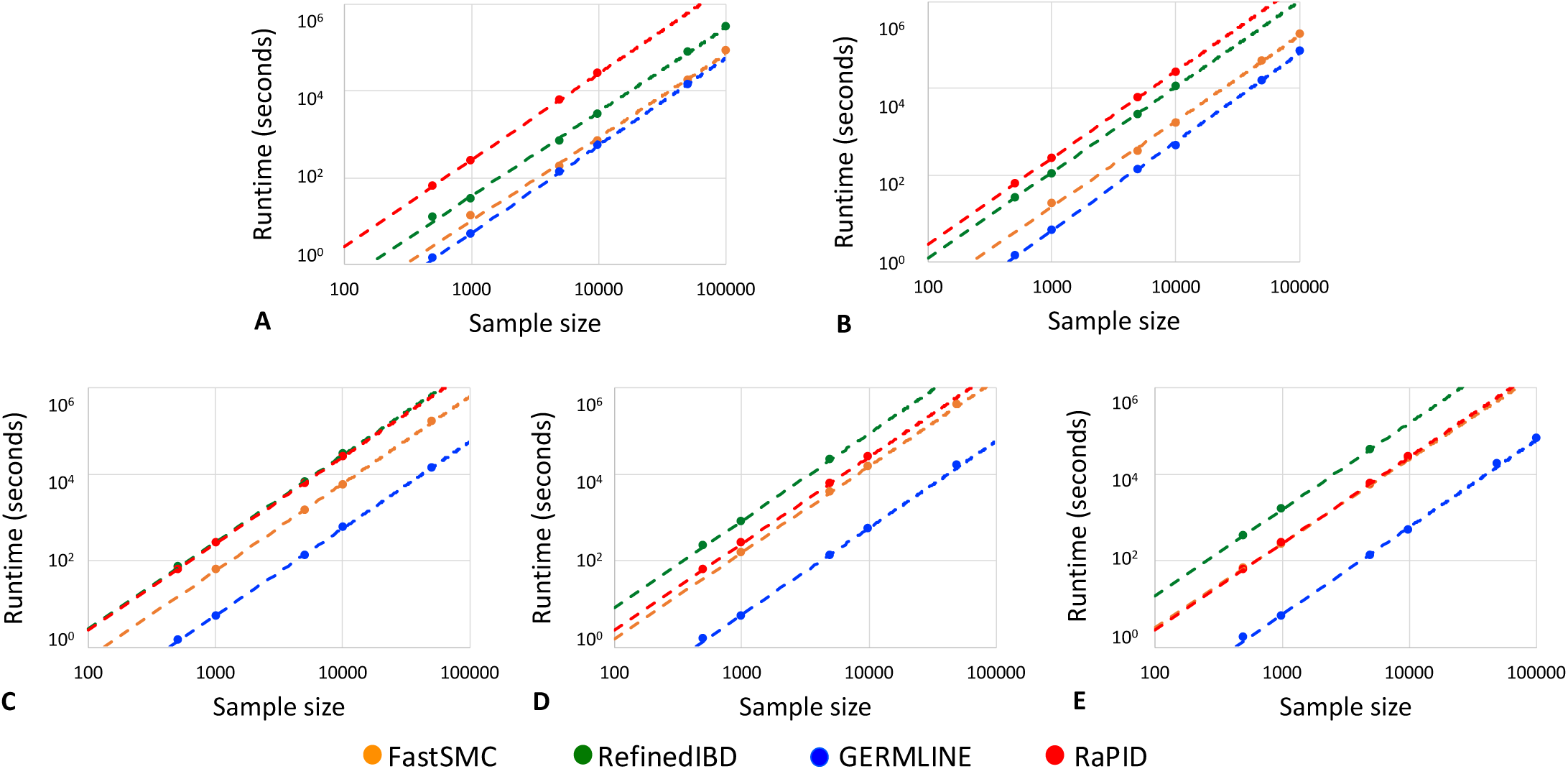
Running time evaluation for IBD detection at different time scales. Running time (CPU seconds) using chromosome 20 of the UKBB across 7,913 SNPs, for IBD segments detection within the past 25 (**A**), 50 (**B**), 100 (**C**), 150 (**D**) and 200 (**E**) generations. The complete cohort of 487,409 samples from the UKBB was randomly downsampled into batches. Only one thread was used for each method (FastSMC, RefinedIBD, GERMLINE and RaPID) and parameters from grid search were used. Trend lines in logarithmic scale reflect differences in the quadratic components of each algorithm.

**Supplementary Fig. 5.**
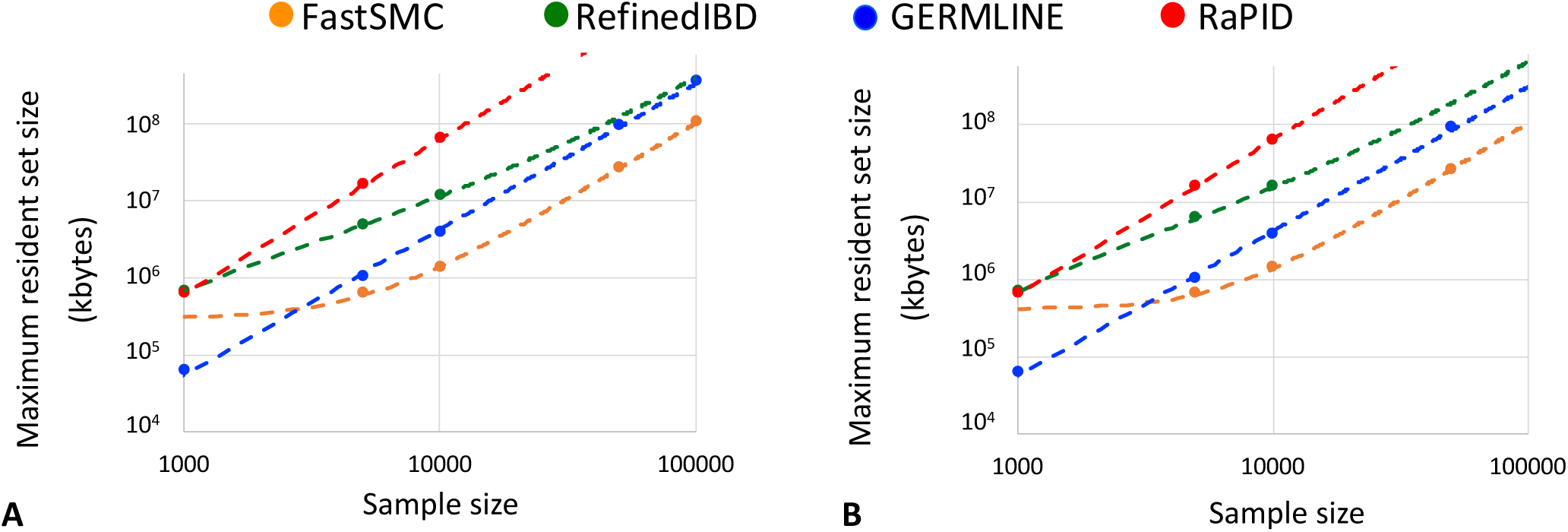
Memory usage for IBD detection at different time scales. Memory usage in kilobytes of FastSMC, RefinedIBD, GERMLINE and RaPID for IBD detection within the past 50 (**A**) and 100 (**B**) generations, with optimal parameter values from the grid search across 7,913 SNPs (chromosome 20). The complete cohort of 487,409 samples from the UKBB was randomly downsampled into batches.

**Supplementary Fig. 6.**
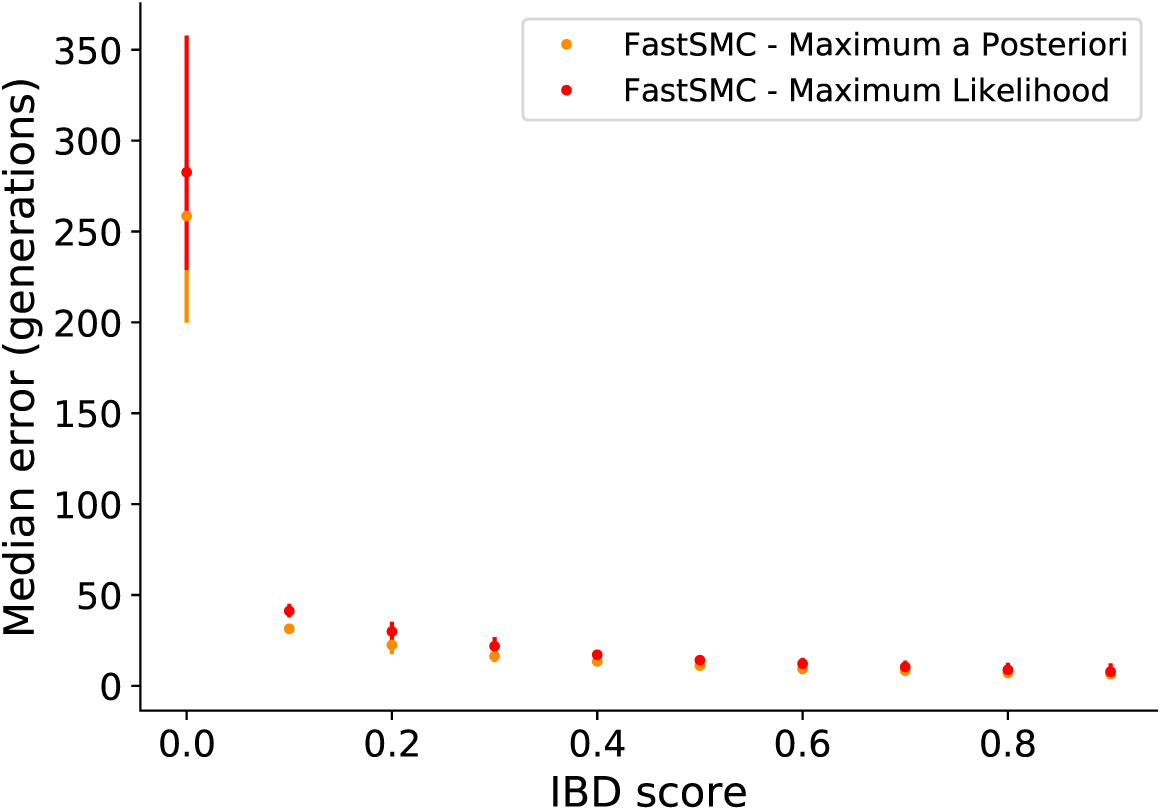
MAP and MLE age estimates comparison. Absolute median error in age estimation in FastSMC for both the MAP and the MLE estimates, at different IBD score thresholds. Vertical lines represent 95% confidence intervals over 10 different simulations (each of them consisting of a 30Mb chromosome under European demographic history model, recombination rates from a human chromosome 2 and SNP ascertainment matching UKBB allele frequencies).

**Supplementary Fig. 7.**
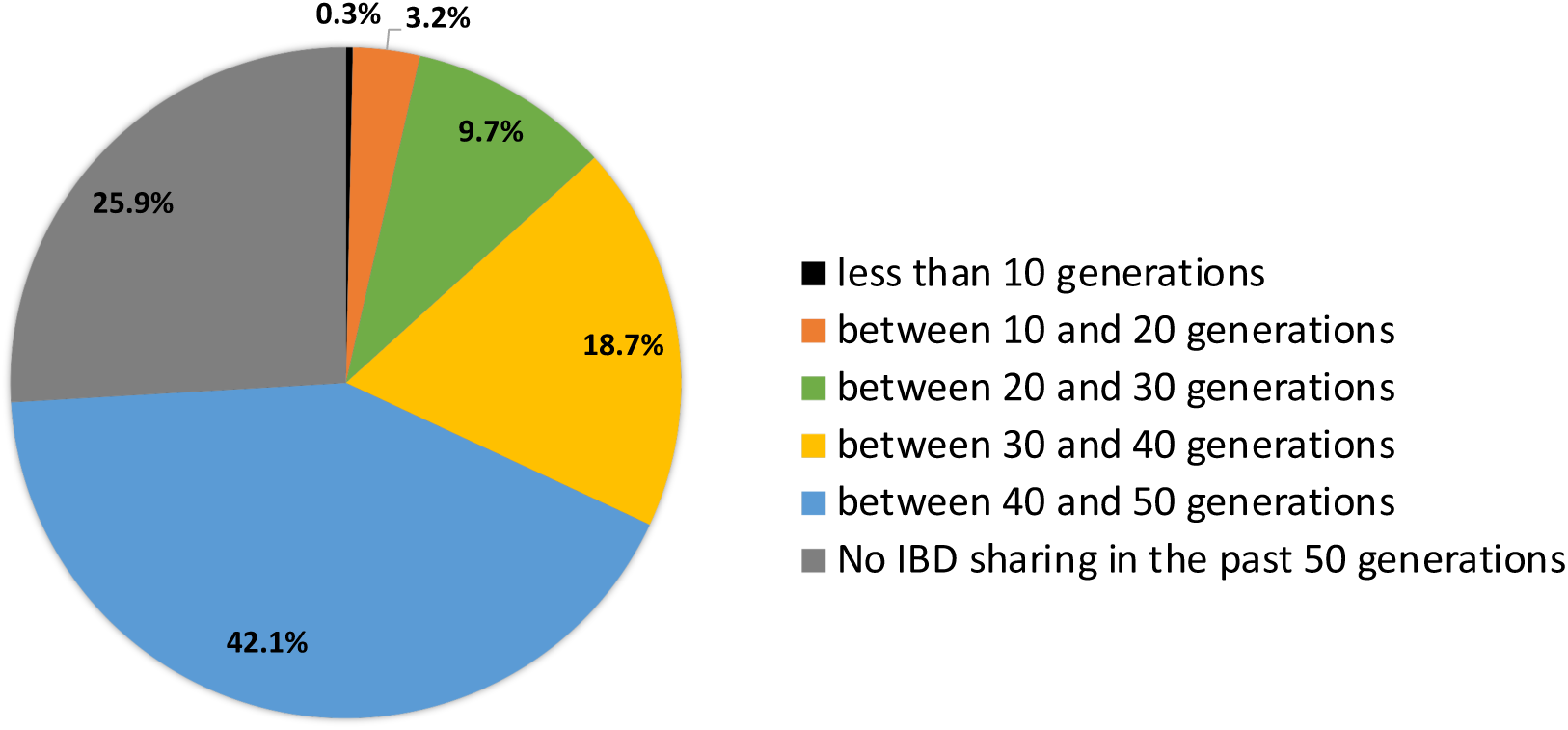
IBD sharing among pairs of samples in the UK Biobank dataset. For each pair of individuals (out of 118,783,522,936 total pairs) in the UK Biobank cohort (487,409 diploid samples) we determined the most recent estimated shared segment, which provides an upper bound for the pair’s genealogical relationship. 74% of all pairs of British individuals were estimated to share common ancestry at some point within the past 50 generations.

**Supplementary Fig. 8.**
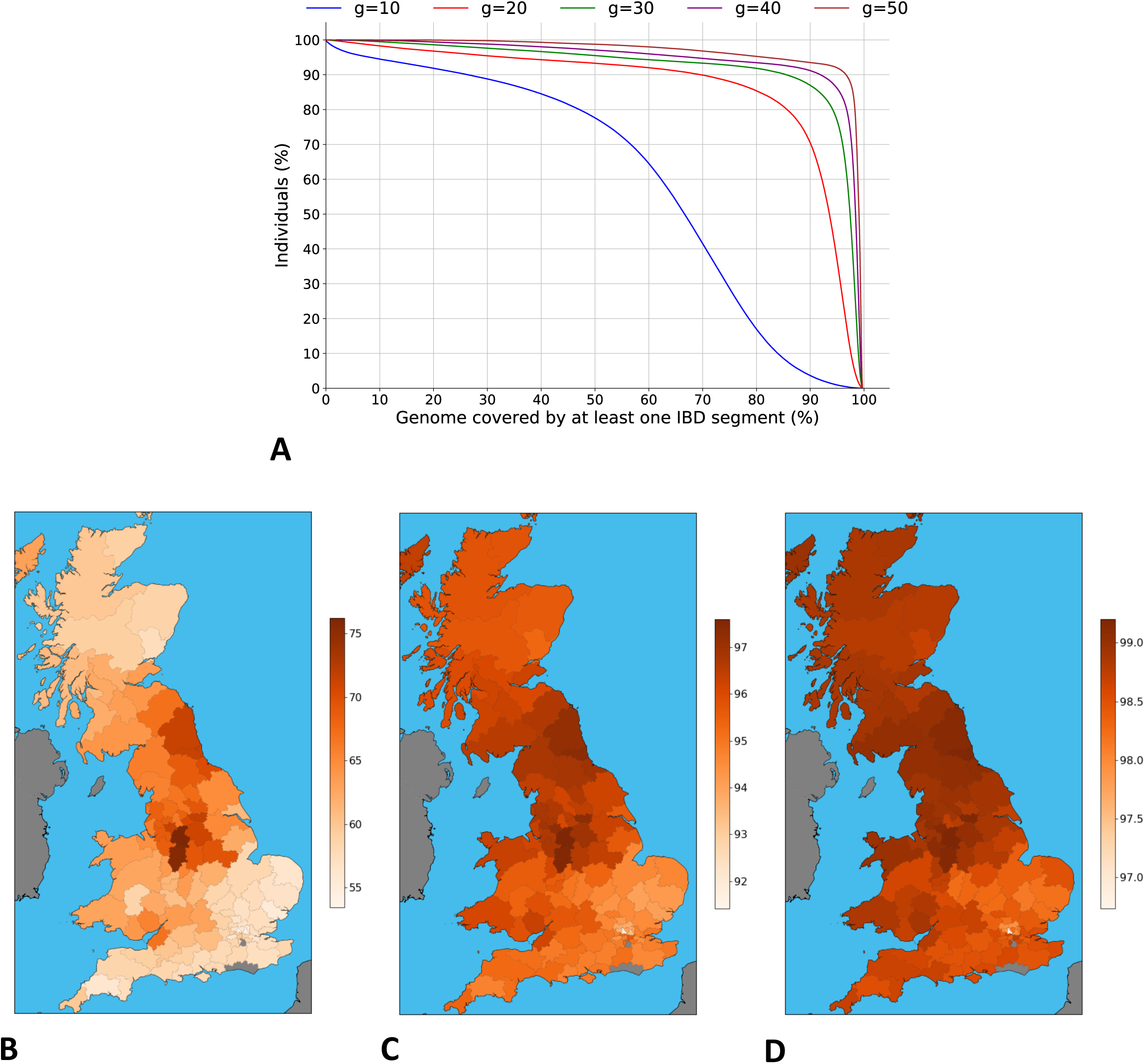
Fraction of genome covered by IBD segments in the UK Biobank dataset. A. Fraction of genome covered by at least one IBD segment (%) for 487,409 samples from the UK Biobank cohort within the past 10, 20, 30, 40 and 50 generations. **B, C, D.** Average fraction of genome covered by at least one IBD segment (%) within UK postcodes for 432,968 samples with known geographic coordinats from the UK Biobank cohort in the past 10 **(B)**, 30 **(C)** and 50 **(D)** generations. Regions with no data available are coloured in gray.

**Supplementary Fig. 9.**
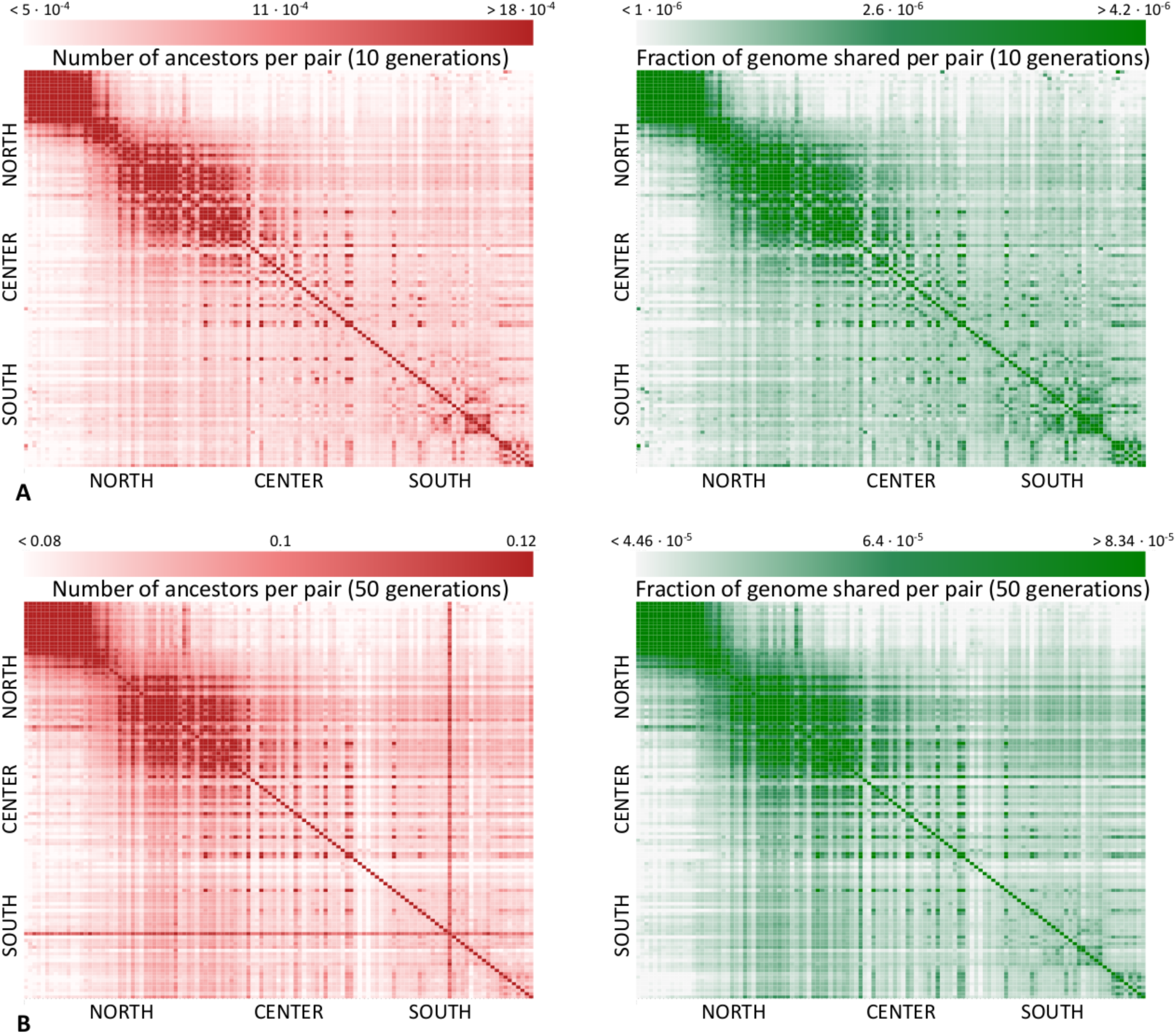
IBD sharing in the past 10 and 50 generations across UK postcodes in the UK Biobank dataset. Average number of IBD segments **(left)** and fraction of genome **(right)** shared per pair of individuals across all pair of postcodes in the UK (excluding Northern Ireland) in the past 10 (**A**) and 50 (**B**) generations. Segments with IBD score smaller than 0.4 were excluded, resulting approximately in recall of 0.6 and precision of 0.8 based on simulations. The red cross appearing in the South of the UK in **B** corresponds to Sutton postcode area (covering south-west London), a cosmopolitan region where samples share many ancestors but small fraction of genome with the rest of the country, revealing deep ancestral ties (i.e short IBD segments) throughout the UK.

**Supplementary Fig. 10.**
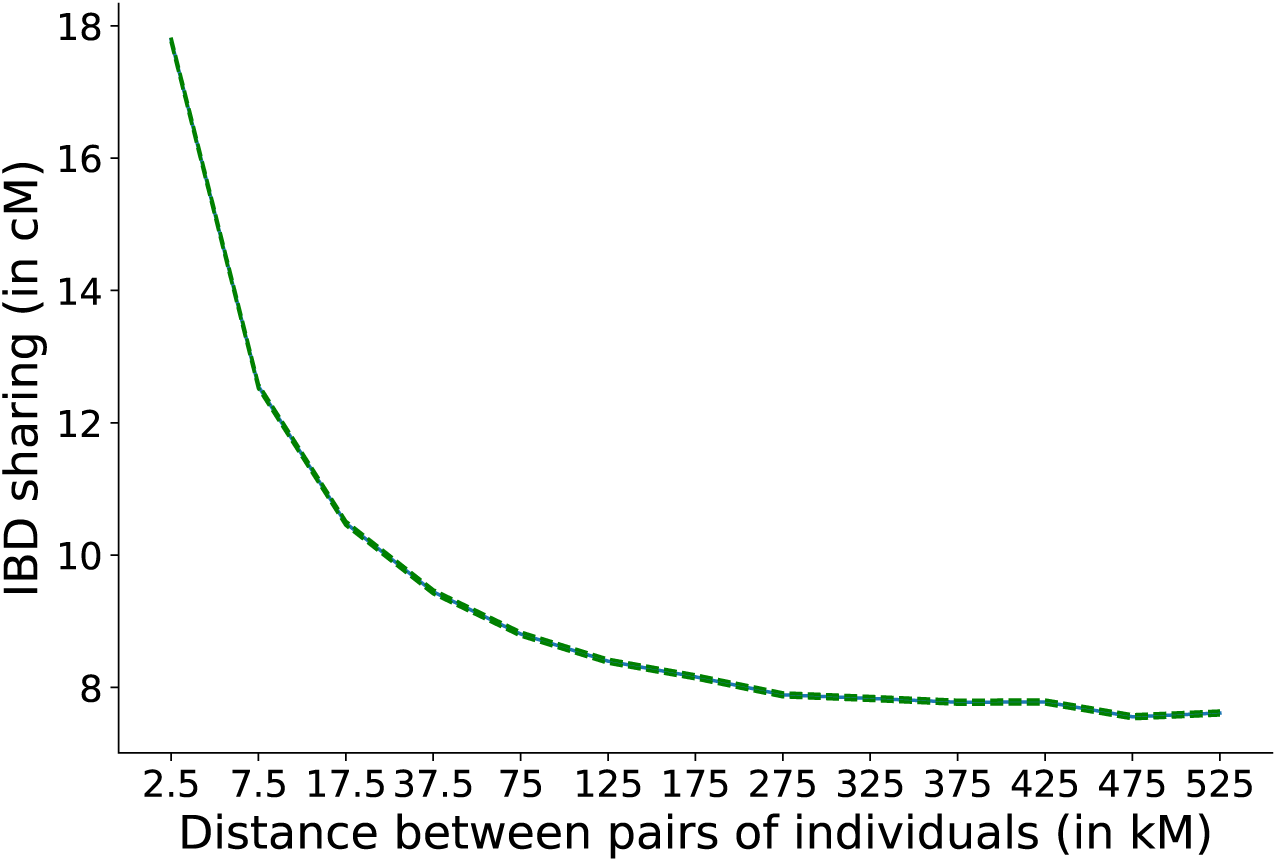
IBD sharing and geographic distance. We randomly sampled 5,000,000 pairs from the UKBB cohort and computed the IBD sharing in cM (i.e sum of IBD segments lengths along the genome within the past 10 generations) for each of them. We partitioned these pairs depending on the distance in kilometers (kM) in the birth locations of the two individuals (less than 5kM, between 5 and 10kM, between 10 and 25kM, between 25 and 50kM, and then every 50kM up to 500kM). The blue line represents the average IBD sharing across all pairs of random samples, and the green trend lines correspond to the standard error of the mean across all pairs (assuming pairs of individuals are independent, which is approximately true when looking at sharing of very recent segments). We observe strong correlation between genetic and geographic distance, demonstrating that close relatives tend to be geographically clustered.

**Supplementary Fig. 11.**
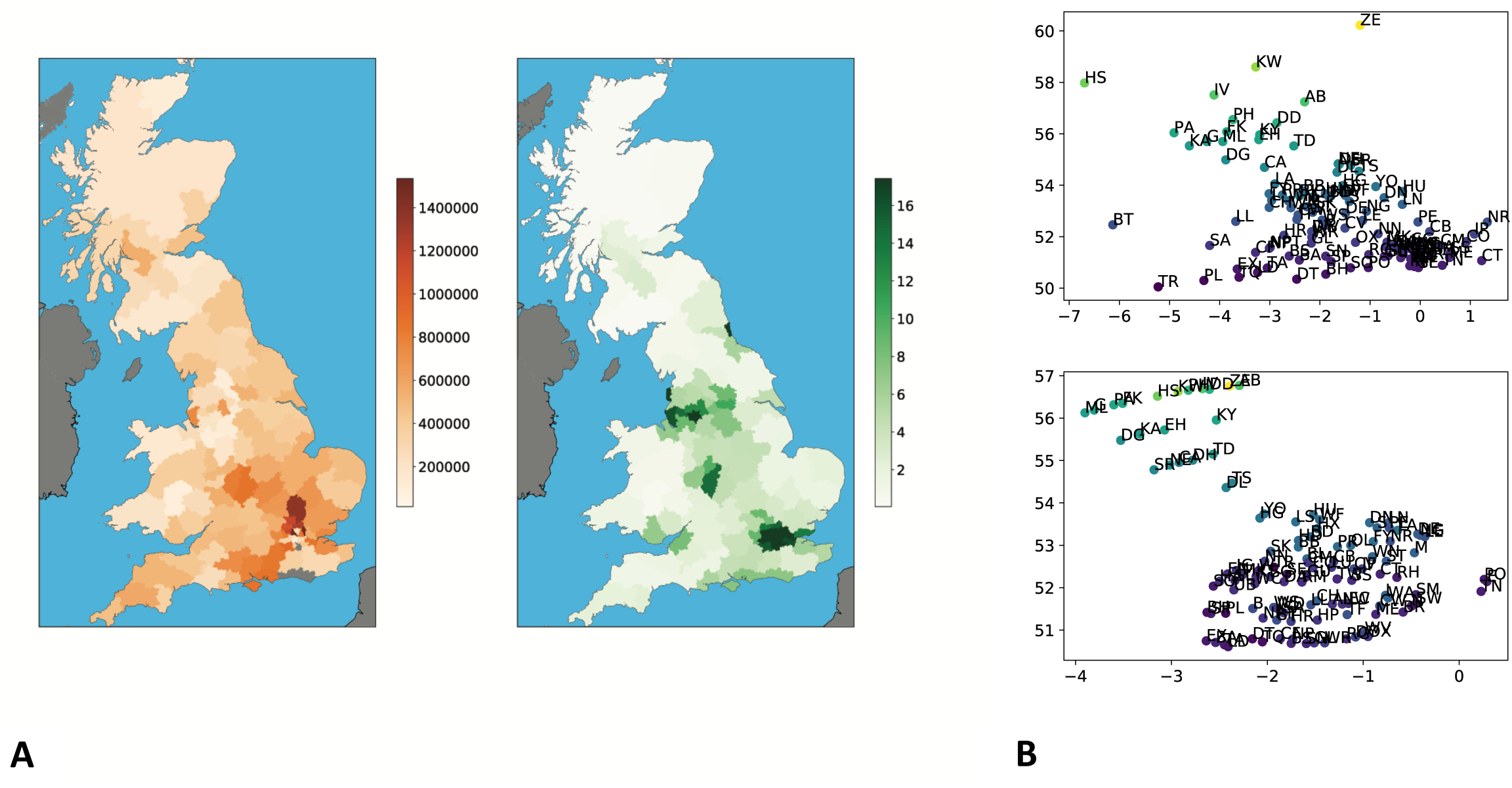
Estimation of effective size and reconstruction of physical distances with IBD sharing. **A.** Recent effective population size estimation based on IBD sharing in the past 10 generations (**left**) and 2011 census population size (number of persons per hectare) within postcodes (**right**). Census population data comes from Nomis web-based dataset for England and Wales, and from the Data Warehouse for Scotland (data only available on the area level, we computed estimates for postcodes). Regions with no data available are coloured in gray. **B.** Real physical distances between postcodes (**top**) and isomap projections using IBD sharing within the past 600 years (**bottom**), x-axis is longitude and y-axis is latitude. The isomap projection was obtained on the IBD sharing postcode dissimilarity matrix and by considering 8 nearest neighbours. We then applied affine transformations (rotation, translation and scaling) to minimize the root mean squared of the reconstruction error (RMSE) (74). The RMSE obtained is 179 km, 95% CI=[163,196].

**Supplementary Fig. 12.**
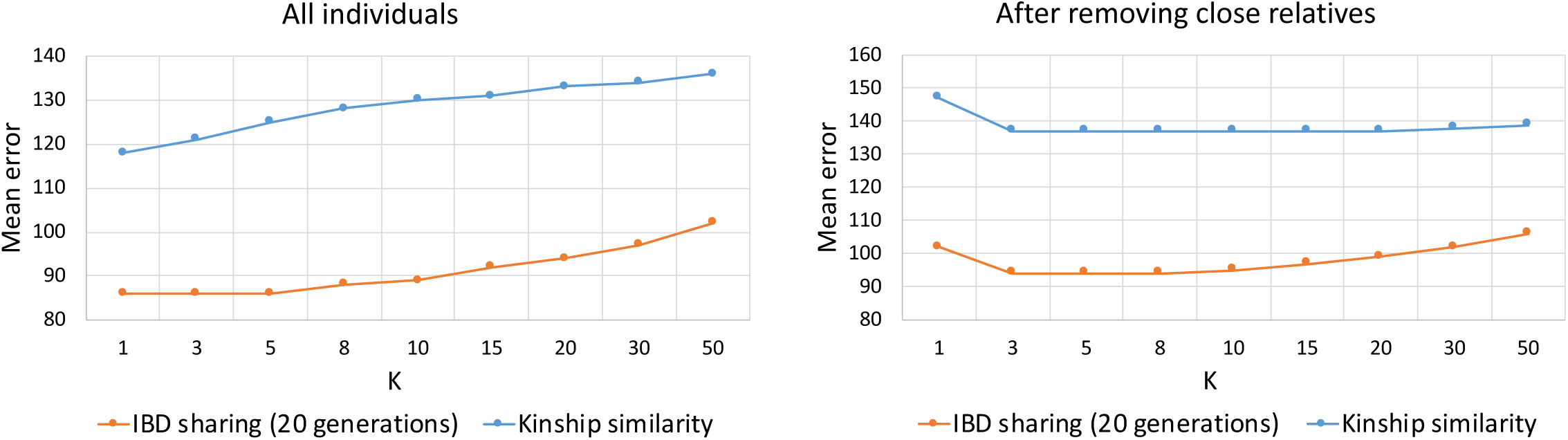
Fine-tuning of the K-nearest neighbours algorithm. **(left)** Average error when predicting birth location of 10,000 random samples from the UKBB, applying the K-nearest-neighbours algorithm while varying the value of the parameter *K*, using both IBD sharing among individuals within the past 20 generations and kinship similarity. **(right)** Average error of 8,226 random samples from the UKBB, after excluding close relatives (≤3rd degree relatives), using the same procedure.

**Supplementary Fig. 13.**
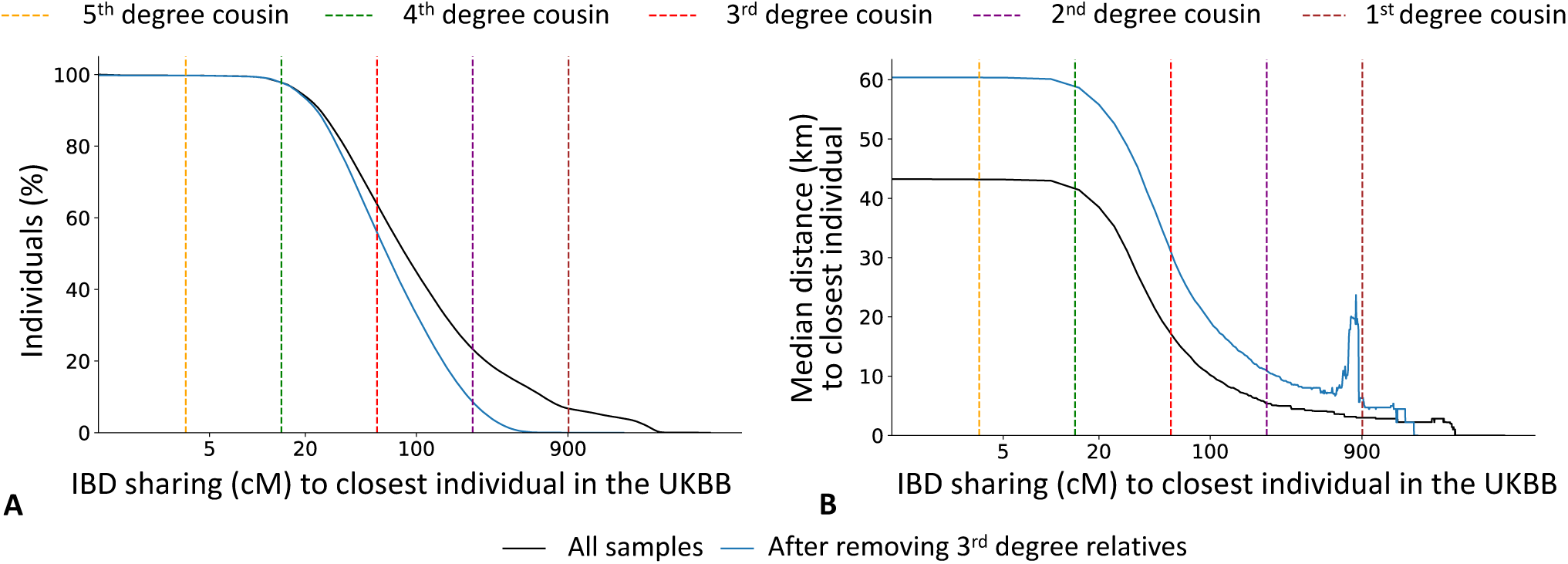
Genetic relatedness and geographic distances in the UK Biobank dataset, before and after removing related individuals. For each UK Biobank sample with available geographic data, we detected the individual sharing the largest total amount (in cM) of genome IBD within the past 10 generations (referred to as “closest individual”). **A.** For each value *x* of total shared genome (in cM) on the X-axis, we report the percentage of UK Biobank samples (Y-axis) that share *x* or more with their closest individual. **B.** For each value *x* of total shared genome (in cM) on the X-axis, we report the median distance (km, computed every 10 cM) for all pairs of (sample, closest individual) who shared at least *x*. Vertical dashed lines indicate the expected value of the total IBD sharing for *k*-th degree cousins, computed using 2*G*(1*/*2)^2(*k*+1)^, where *G* = 7247.14 is the total diploid genome size (in cM) and *k* represents the degree of cousin relationship (e.g. *k* = 2 for second degree cousins, separated by 2(*k* + 1) generations) (10). We show results obtained using either all individuals with available geographic data (*N* = 432,968; black lines), or all individuals with available geographic data after removing ≤3rd degree relatives (e.g. first degree cousins), detected in (2) using the KING software (72) (*N* = 357,588; blue lines).

**Supplementary Fig. 14.**
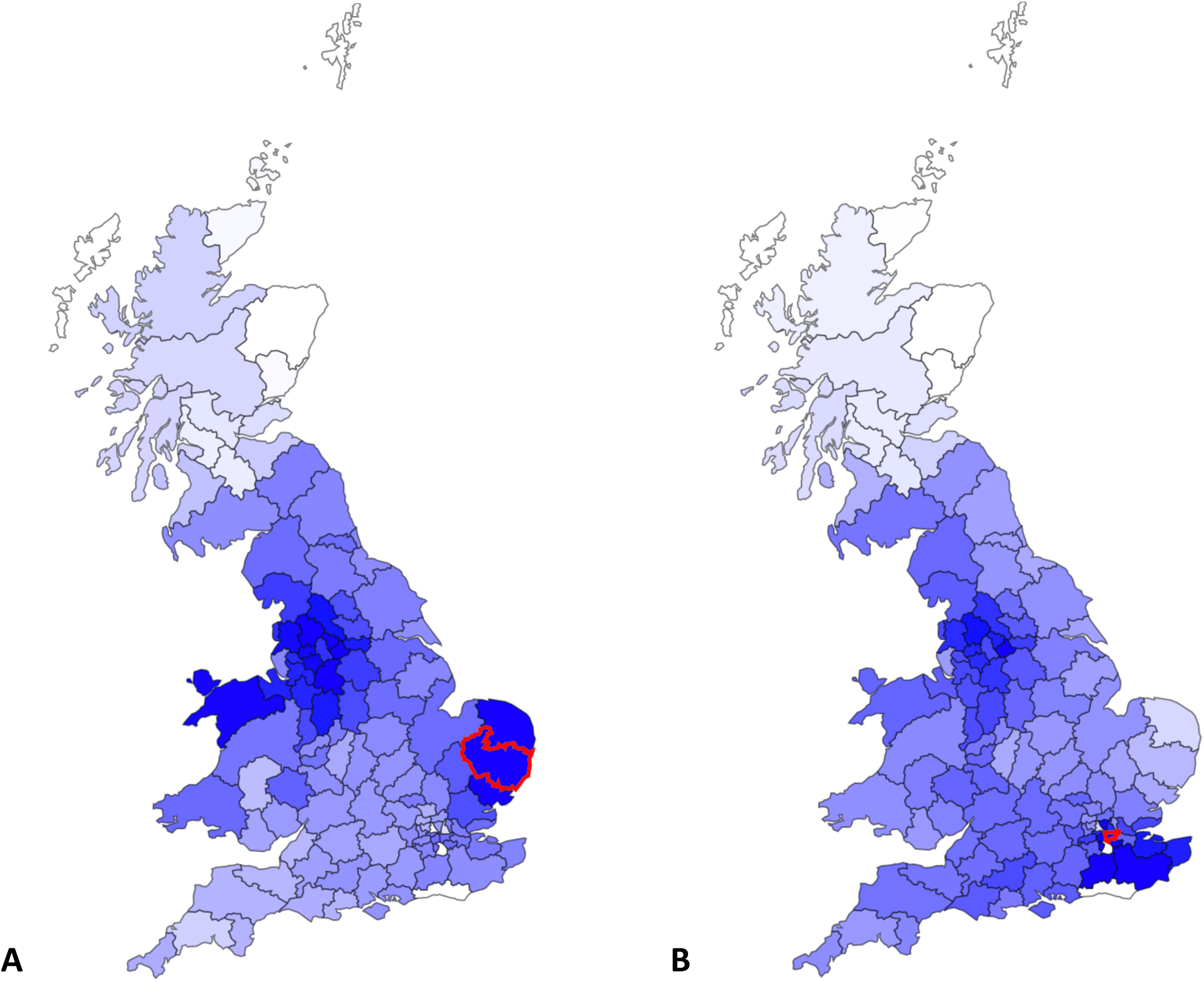
Pervasive IBD sharing with North-West England. Individuals throughout the UK, including individuals from cosmopolitan regions such as London, have deep genetic relationship with modern-day individuals from North-West England, in addition to nearby regions. This figure displays the average fraction of genome shared through IBD segments in the past 1,500 years per pair of individuals between the IP postcode (corresponding to Ipswich, in red) and other UK regions (**A**), and between the SE postcode (corresponding to South East London, in red) and other UK regions (**B**). The darker the color is, the higher the average fraction of genome shared is. More details and an interactive map can be found at https://ukancestrymap.github.io/

**Supplementary Fig. 15.**
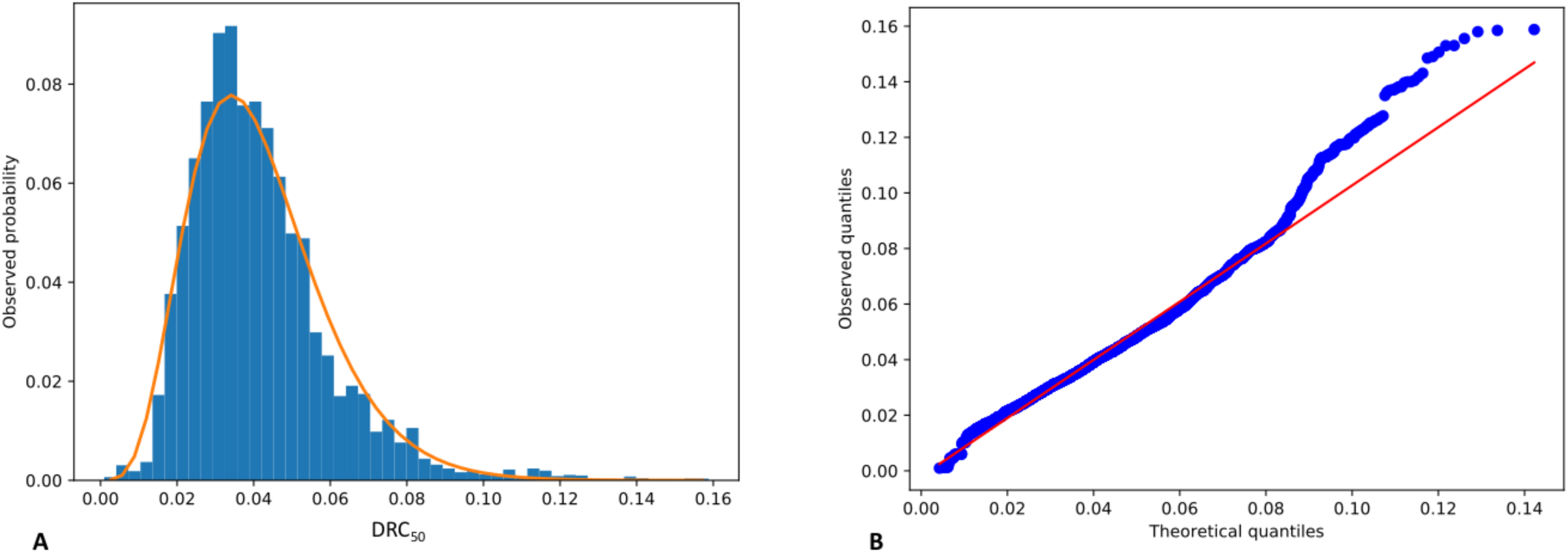
Empirical null model for the DRC_50_ statistic. Empirical null model for detection of recent positive selection, fitted using a Gamma distribution with shape, location and scale parameters (see Methods). **A.** Empirical distribution (in blue) and Gamma-fit (orange curve) for the DRC_50_ statistic in the putative neutral regions of the genome (see URLs) in the UKBB, after excluding significant loci falling within these putative neutral regions (9,165 observations from 0.05 cM windows). **B.** Quantile-quantile plot for the DRC_50_ statistic in the putative neutral regions of the genome (see URLs) in the UKBB, after excluding significant loci falling within these putative neutral regions (9,165 observations from 0.05 cM windows).

**Supplementary Fig. 16.**
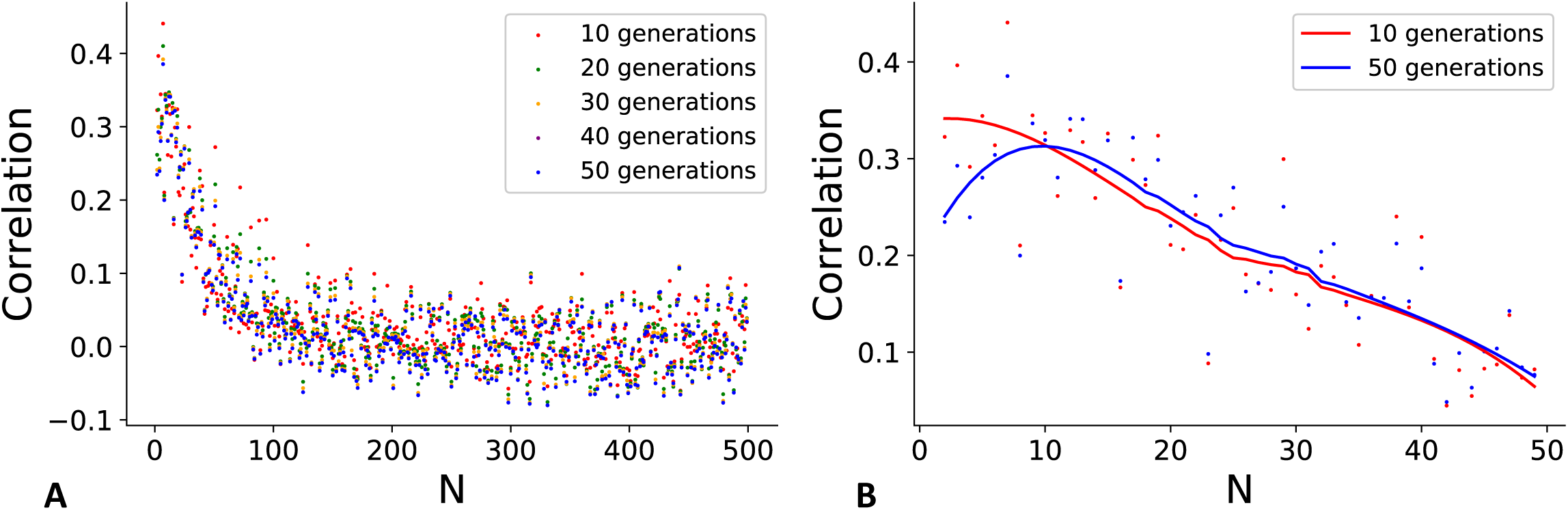
Correlation between IBD sharing and sharing of rare variants. **A.** Correlation between IBD sharing (average number of IBD segments per pair within UK regions in the past 10, 20, 30, 40 and 50 generations in the UK Biobank cohort) and ultra-rare variants sharing (average number of *F*_*N*_ mutations per pair within UK regions in the UK Biobank 50k Exome Sequencing Data Release, for any *N* between 2 and 499). **B.** Non-parametric regression of the correlation between IBD sharing (average number of IBD segments per pair within UK regions in the past 10 and 50 generations in 487,409 samples in the UK Biobank dataset) and ultra-rare variants sharing (average number of *F*_*N*_ mutations per pair within UK regions in the UK Biobank 50k Exome Sequencing Data Release, for values of *N* smaller than 50).

**Supplementary Fig. 17.**
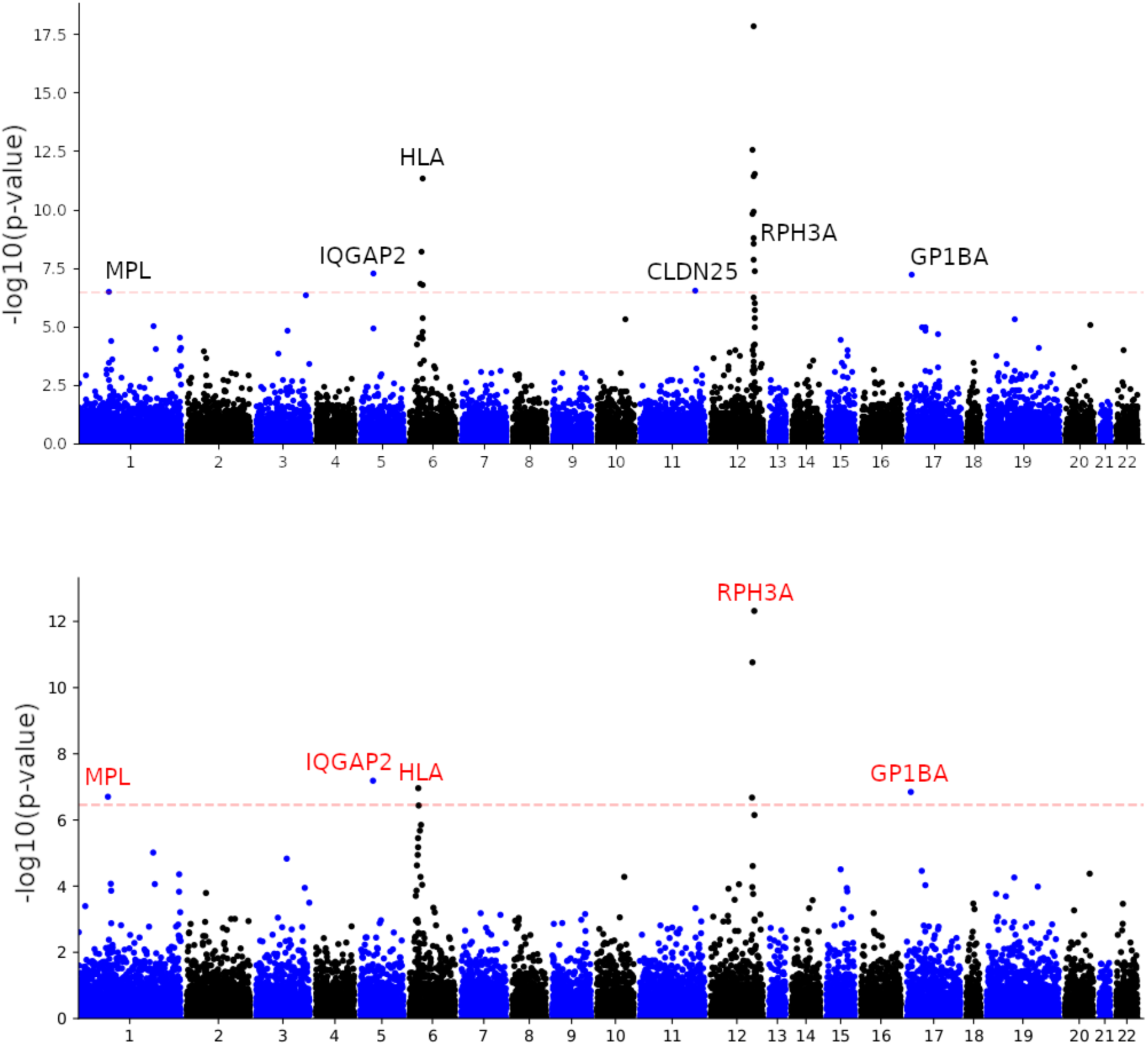
LoF-segment burden exome-wide Manhattan plots for platelet count with and without SNP-adjustment. Labelled genes are exome-wide significant after adjusting for multiple testing (t-test p < 0.05*/*(14,249 × 10) = 3.51 × 10^−7^; dashed red line). We compare results before (**top**) and afer (**bottom**) adjusting for common SNP associations. Both LoF-segment burden analyses used 303,125 UK Biobank samples not included in the exome sequencing cohort. The cluster of genes in chromosome 12 labeled as *RPH3A* (the top association) contains *KCTD10, TCHP* and *RPH3A* and the signal with *CLDN25* was cleared after SNP-adjustment. Red labels in the bottom plot indicate associations that were not detected in our WES-based LoF burden analysis or reported by Van Hout et al.

**Supplementary Fig. 18.**
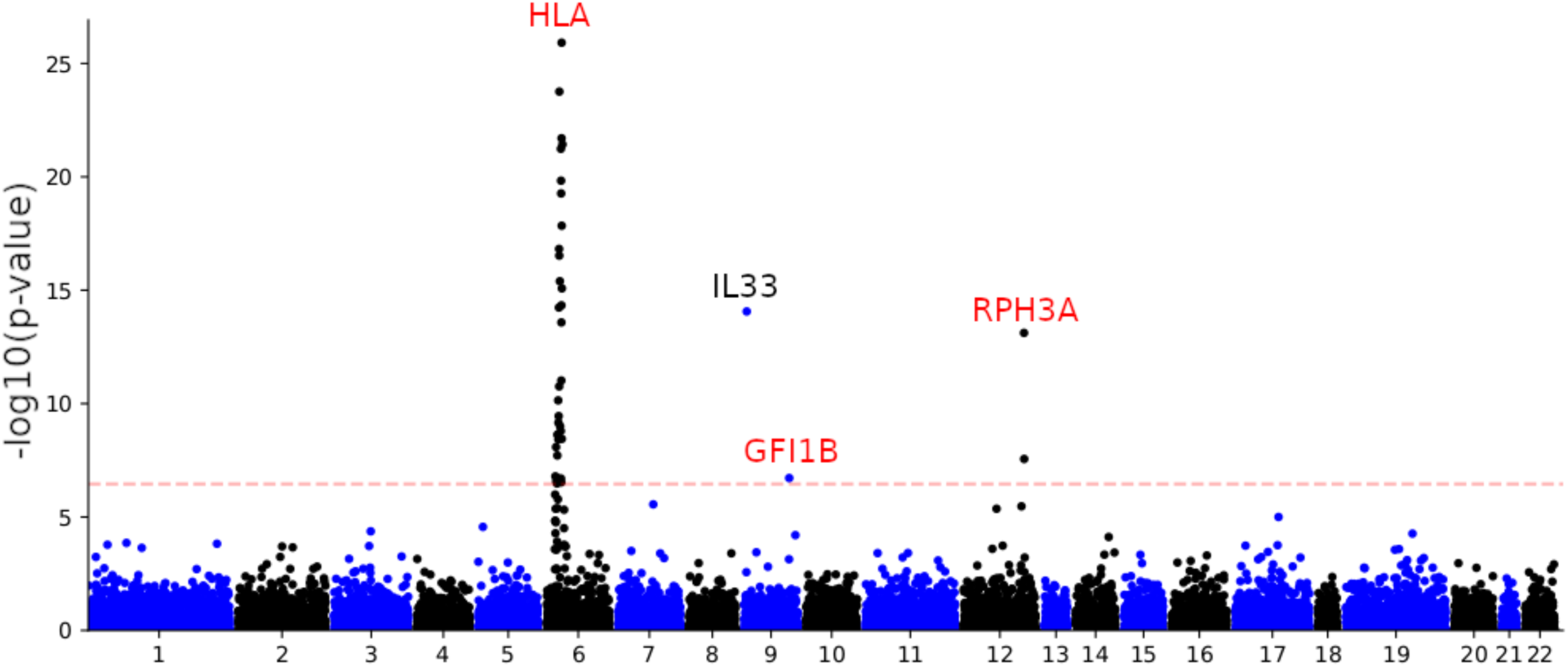
LoF-segment burden exome-wide Manhattan plot for eosinophill count. Labelled genes are exome-wide significant (after adjusting for multiple testing, t-test p-value < 0.05*/*(14,249 × 10) = 3.51 × 10^−7^; dashed red line). The LoF-segment burden analysis (with SNP adjustment) used 303,125 UK Biobank samples not included in the exome sequencing cohort. We identified one locus previously reported by Van Hout et al. (black label), and additional loci on chromosomes 6,9 and 12 (labels in red). Table 2 reports the list of genes within the gene cluster labeled as HLA.

**Supplementary Fig. 19.**
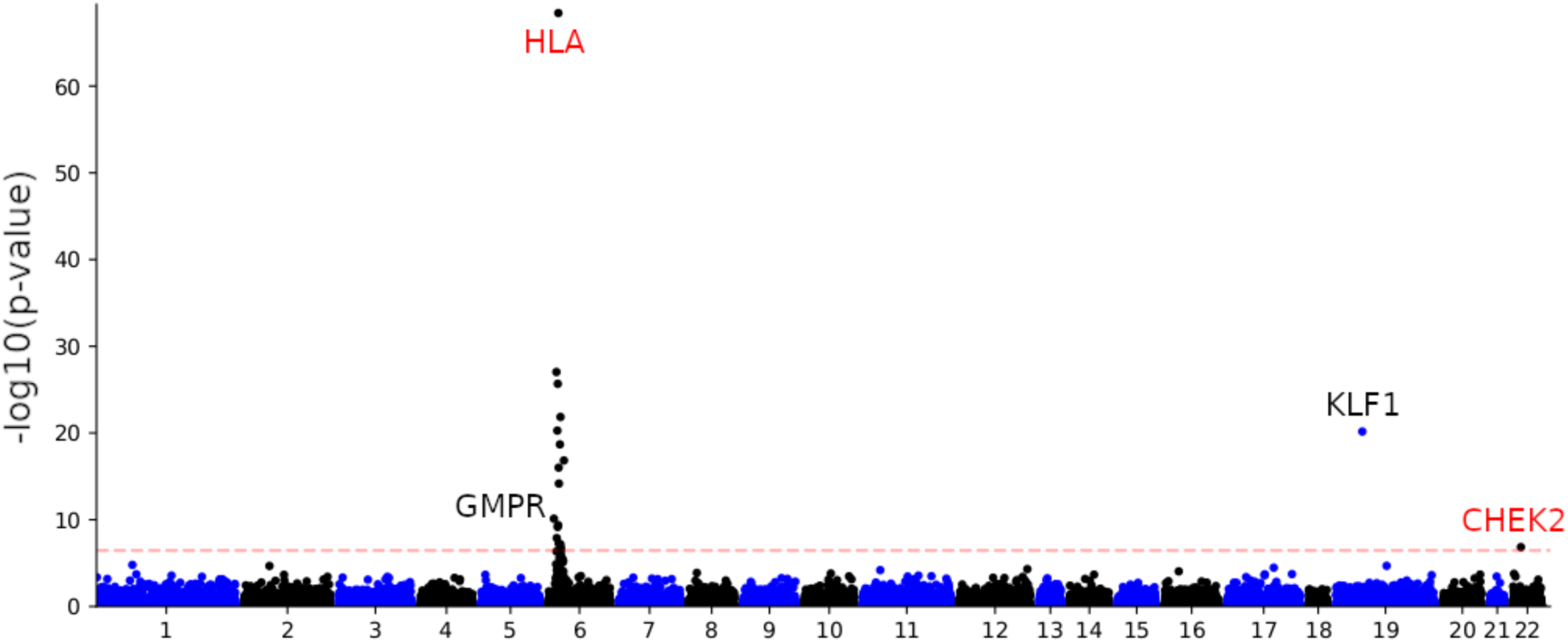
LoF-segment burden exome-wide Manhattan plot for mean corpuscular haemoglobin. Labelled genes are exome-wide significant (after adjusting for multiple testing, t-test p-value < 0.05*/*(14,249 × 10) = 3.51 × 10^−7^; dashed red line). The LoF-segment burden analysis (with SNP adjustment) used 303,125 UK Biobank samples not included in the exome sequencing cohort. We identified two loci previously reported by Van Hout et al. (45), *KLF1* and *GMPR* (gene labels in black), and two novel associations at HLA and *CHEK2* (labels in red).

**Supplementary Fig. 20.**
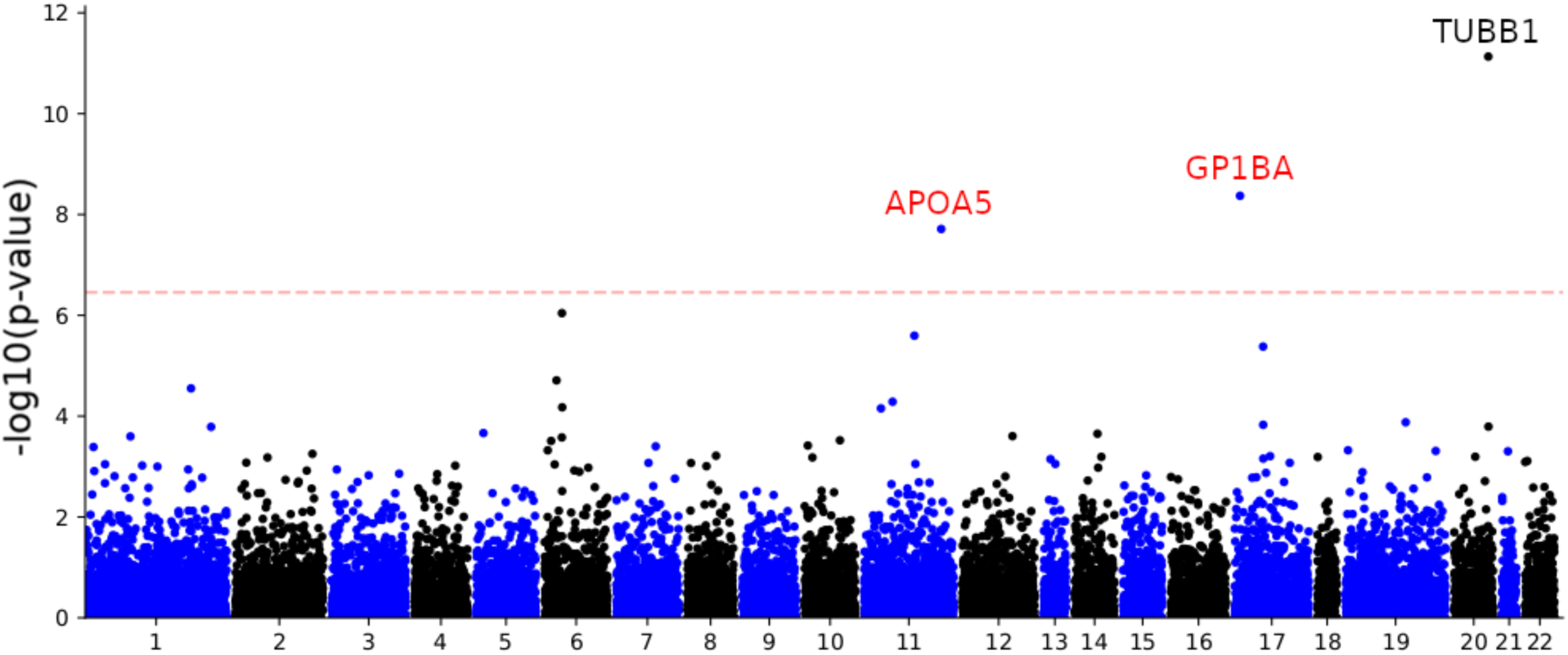
LoF-segment burden exome-wide Manhattan plot for platelet distribution width. Labelled genes are exome-wide significant (after adjusting for multiple testing, t-test p-value < 0.05*/*(14,249 × 10) = 3.51 × 10^−7^; dashed red line). The LoF-segment burden analysis (with SNP adjustment) used 303,125 UK Biobank samples not included in the exome sequencing cohort. We identified one locus previously reported by Van Hout et al. (45), *TUBB1* (black label), and two additional genes, *APOA5* and *GP1BA* (labels in red).

**Supplementary Fig. 21.**
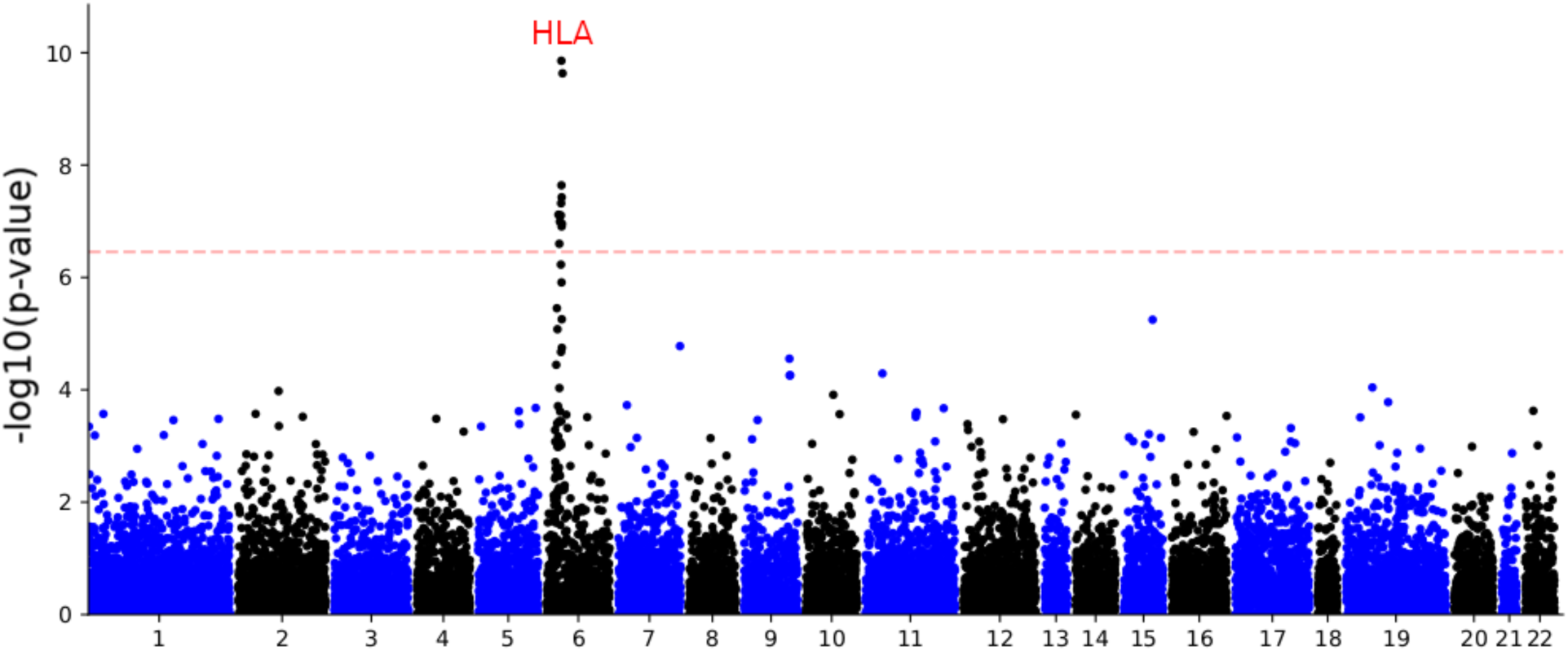
LoF-segment burden exome-wide Manhattan plot for red blood cell count. Labelled genes are exome-wide significant (after adjusting for multiple testing, t-test p-value < 0.05*/*(14,249 × 10) = 3.51 × 10^−7^; dashed red line). The LoF-segment burden analysis (with SNP adjustment) used 303,125 UK Biobank samples not included in the exome sequencing cohort. We detected one novel association at the HLA locus which was not detected by either of the WES-burden tests.

**Supplementary Fig. 22.**
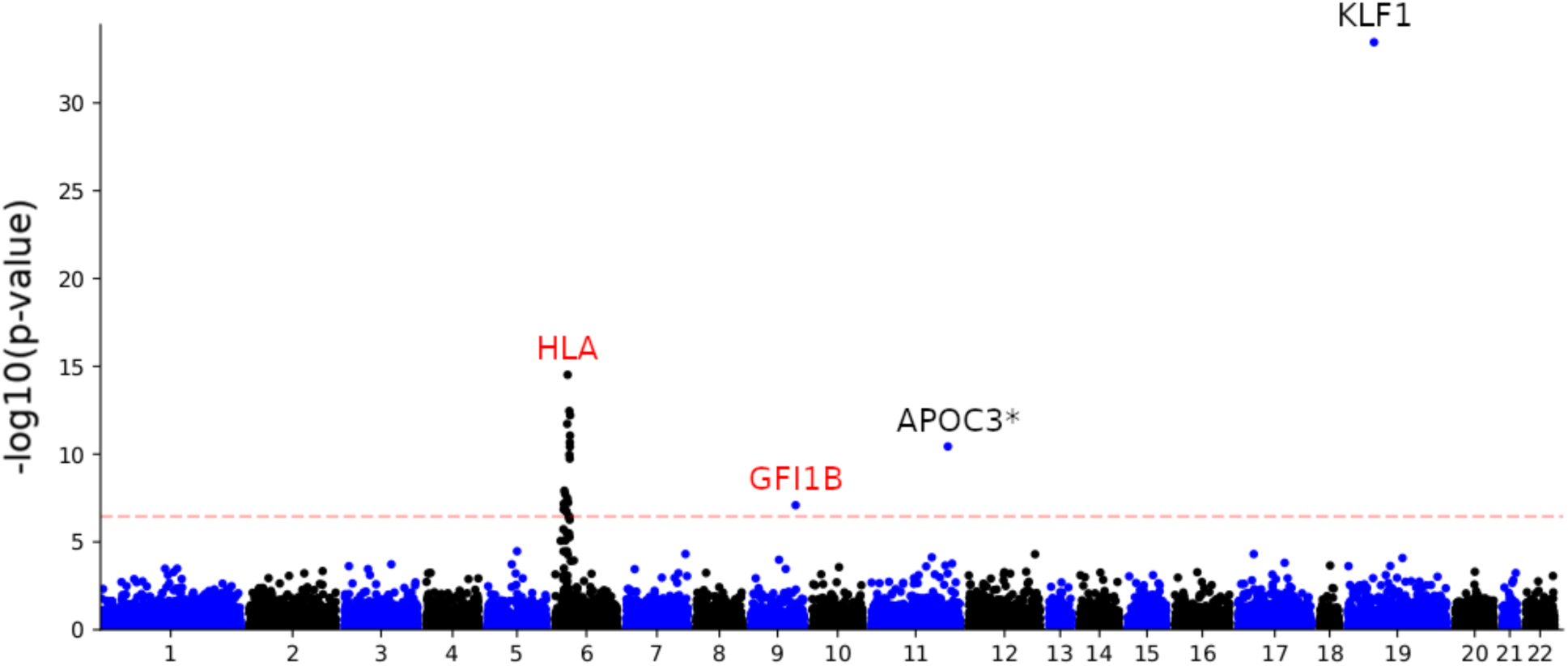
LoF-segment burden exome-wide Manhattan plot for red blood cell distribution width. Labelled genes are exome-wide significant (after adjusting for multiple testing, t-test p-value < 0.05*/*(14,249 × 10) = 3.51 × 10^−7^; dashed red line). The LoF-segment burden analysis (with SNP adjustment) used 303,125 UK Biobank samples not included in the exome sequencing cohort. We identified two previously-reported loci (black labels), *KLF1* (detected by Van Hout et al. (45)) and *APOC3* (detected by our WES burden analysis), along with two additional (labels in red).

**Supplementary Fig. 23.**
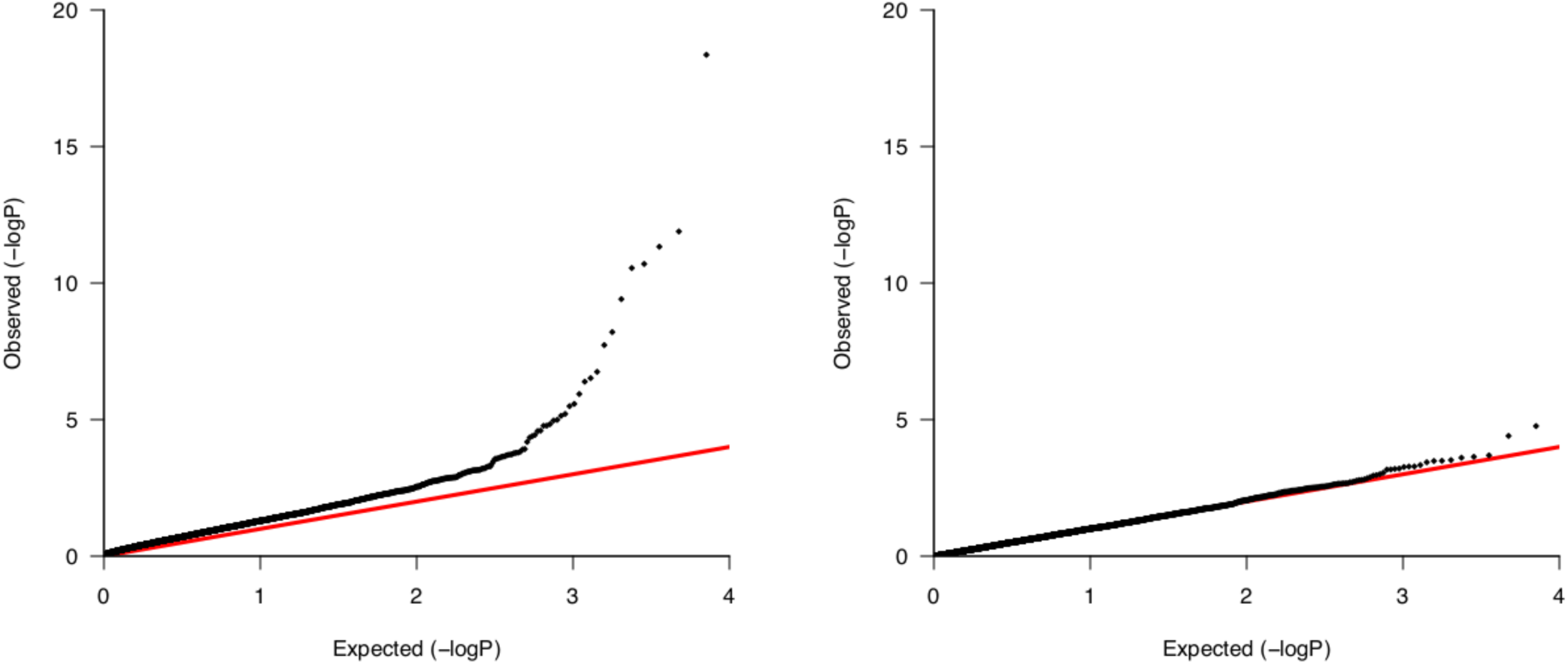
Quantile-quantile plots for LoF-segment burden association. Quantile-quantile plots for mean platelet (thrombocyte) volume **(left)** and for the same trait, but with randomly permuted phenotype values **(right)**. We observe no signal in the permuted phenotype analysis, suggesting a well-calibrated test. The genomic inflation factor for the (SNP-adjusted) LoF-segment burden test, calculated as the the ratio between the observed median chi-squared association statistic and the median chi-squared association statistic expected under the null, is 1.982 (similar values were observed for the other traits). Values larger than 1 may be caused by pervasive polygenicity (75) and are also observed in analyses of common variants (1.492 for this trait using summary statistics from (73)) and by population stratification remaining after correcting for principal components (67) (see Discussion).

## SUPPLEMENTARY TABLES

**Supplementary Table 1.** Optimal parameters from the grid search for IBD detection methods. The accuracy of each method (FastSMC, GERMLINE, RaPID and RefinedIBD) was optimised at different time scale (25, 50, 100, 150 and 200 generations) on a simulated dataset of 300 haploid individuals from a European demographic model and a region of 30 Mb from chromosome 2 (see Methods). FastSMC has two main parameters that we tuned: a time threshold (**t**) to indicate how deep in time the user wants to detect common ancestry, and a minimum length parameter (in cM) (**length**) for the IBD segments. The time threshold parameter was always set to be the same as the time threshold used for the benchmarking. We tuned three parameters in GERMLINE: the minimum length of IBD segments (in cM) (**min**, optimized over [0.001, 0.01, 0.1, 0.5, 0.75, 1, 1.5, 3, 5]), the numbers of bits (SNPs) in each window (**bits**, optimized over [32, 64, 128, 256]), the minimum number of mismatches allowed in each of them (**err**, optimized over [0, 2, 5, 10, 20]). RaPID has four major parameters: the minimum IBD segments length (in cM) (**l**, optimized over [0.001, 0.01, 0.1, 0.5, 1, 1.5, 3, 5]), the number of iterations for the PBWT algorithm (**r**, optimized over [1, 5, 10, 40, 70]), the minimum number of successes (**s**, optimized over [1, 5, 8, 10, 20, 35, 40, 50, 70]) and the window size (in SNPs) (**w**, optimized over [1, 3, 5, 10, 15, 20, 50, 85, 110, 150]). RefinedIBD’s parameters include a minimum length (in cM) (**length**, optimized over [0.001, 0.01, 0.1, 0.5, 0.75, 1, 1.5, 3, 5]) and a minimum LOD score (proxy for quality) (**lod**, optimized over [0.01, 0.1, 1, 3, 5]). We used default values for all parameters not mentioned here.

| Time |  | 25 | 50 | 100 | 150 | 200 |
| --- | --- | --- | --- | --- | --- | --- |
| Method |  |  |  |  |  |  |
| FastSMC | **t** | 25 | 50 | 100 | 150 | 200 |
|  | **length** | 1.5 | 1 | 0.5 | 0.25 | 0.1 |
| GERMLINE | **min** | 0.001 | 0.001 | 0.001 | 0.001 | 0.001 |
|  | **bits** | 64 | 64 | 64 | 64 | 64 |
|  | **err** | 0 | 0 | 0 | 0 | 0 |
| RaPID | **l** | 0.5 | 0.5 | 0.5 | 0.5 | 0.5 |
|  | **r** | 10 | 10 | 10 | 10 | 10 |
|  | **s** | 8 | 8 | 8 | 5 | 5 |
|  | **w** | 3 | 3 | 3 | 3 | 3 |
| RefinedIBD | **length** | 1.5 | 1 | 0.75 | 0.5 | 0.1 |
|  | **lod** | 1 | 1 | 1 | 1 | 1 |

**Supplementary Table 2.**
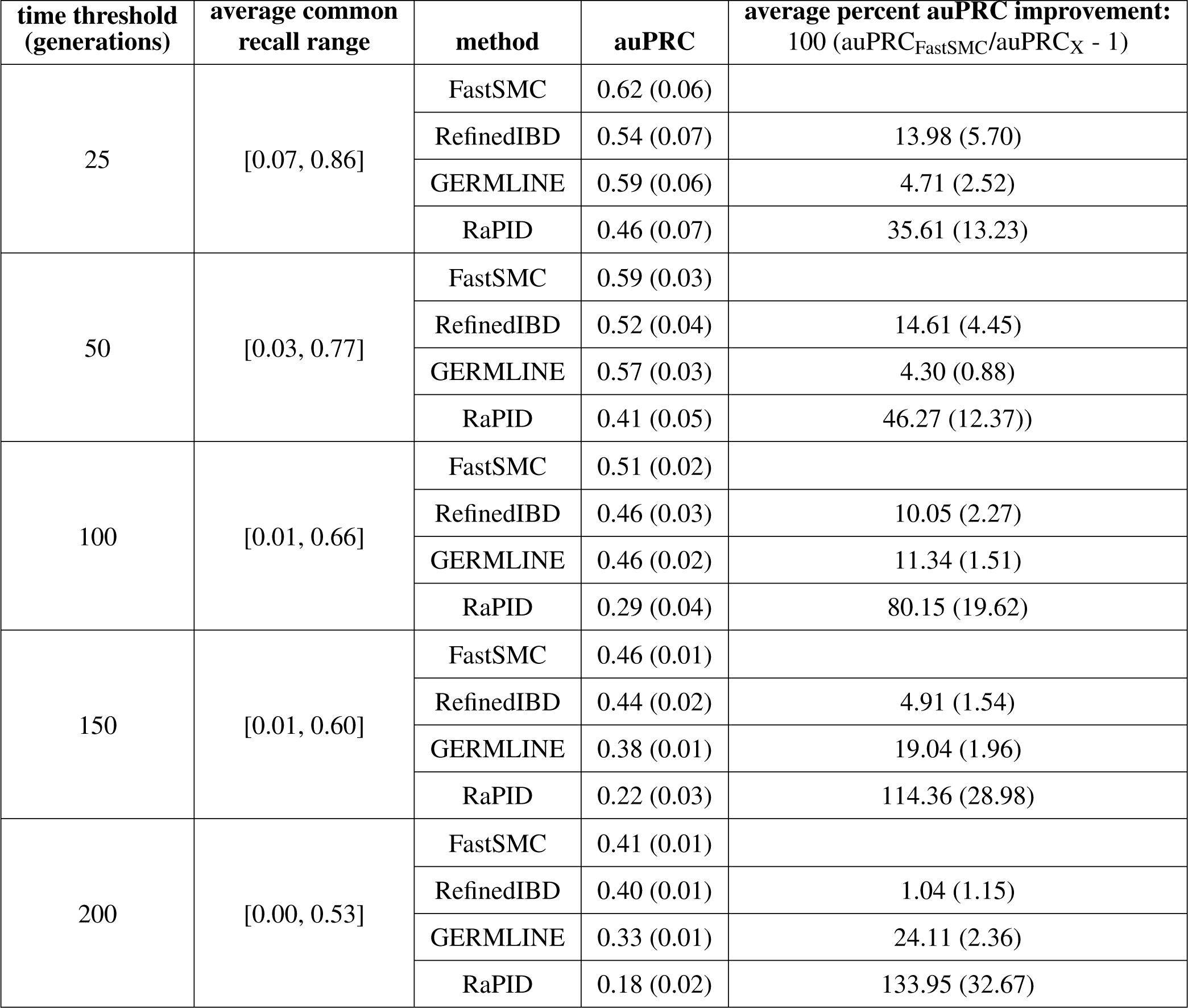
Accuracy measurement. Difference in accuracy between FastSMC and other IBD detection methods within the past 25, 50, 100, 150 and 200 generations. We report the percent improvement for the area under the precision-recall curve (auPRC) of FastSMC over other methods. For all methods, precision can only be estimated within a limited recall range (due mostly to the minimum length parameter) and we only report accuracy measurement on the common recall range. Optimal parameters from fine-tuning were used (see Supplementary Table 1). Accuracy was measured on 10 realistic simulated datasets, all different from the one used for parameters fine-tuning and all consisting of a 30Mb chromosome under European demographic history model for 300 samples, recombination rates from a human chromosome 2 and SNP ascertainment matching UKBB allele frequencies (see Methods). Numbers in round brackets represent standard errors over 10 simulations.

**Supplementary Table 3.** Effects of demographic model misspecification. We simulated 10 batches of 300 haploid samples from the first 30Mb of a human Chromosome 2 and a constant population size of 10, 000 diploid individuals. We ran FastSMC at different time scales, assuming a European demographic model and we reported the auPRC and the average recall range. Numbers in round brackets represent standard errors. Values are very similar to results obtained on samples simulated with a European demographic model (see Supplementary Table 2). Demographic model misspecification does not impact auPRC.

| **time threshold (generations)** | **average recall range** | **auPRC** |
| --- | --- | --- |
| 25 | [0.07, 0.84] | 0.62 (0.09) |
| 50 | [0.03, 0.72] | 0.57 (0.05) |
| 100 | [0.01, 0.65] | 0.51 (0.03) |
| 150 | [0.0, 0.59] | 0.45 (0.02) |
| 200 | [0.0, 0.55] | 0.4 (0.01) |

**Supplementary Table 4.**
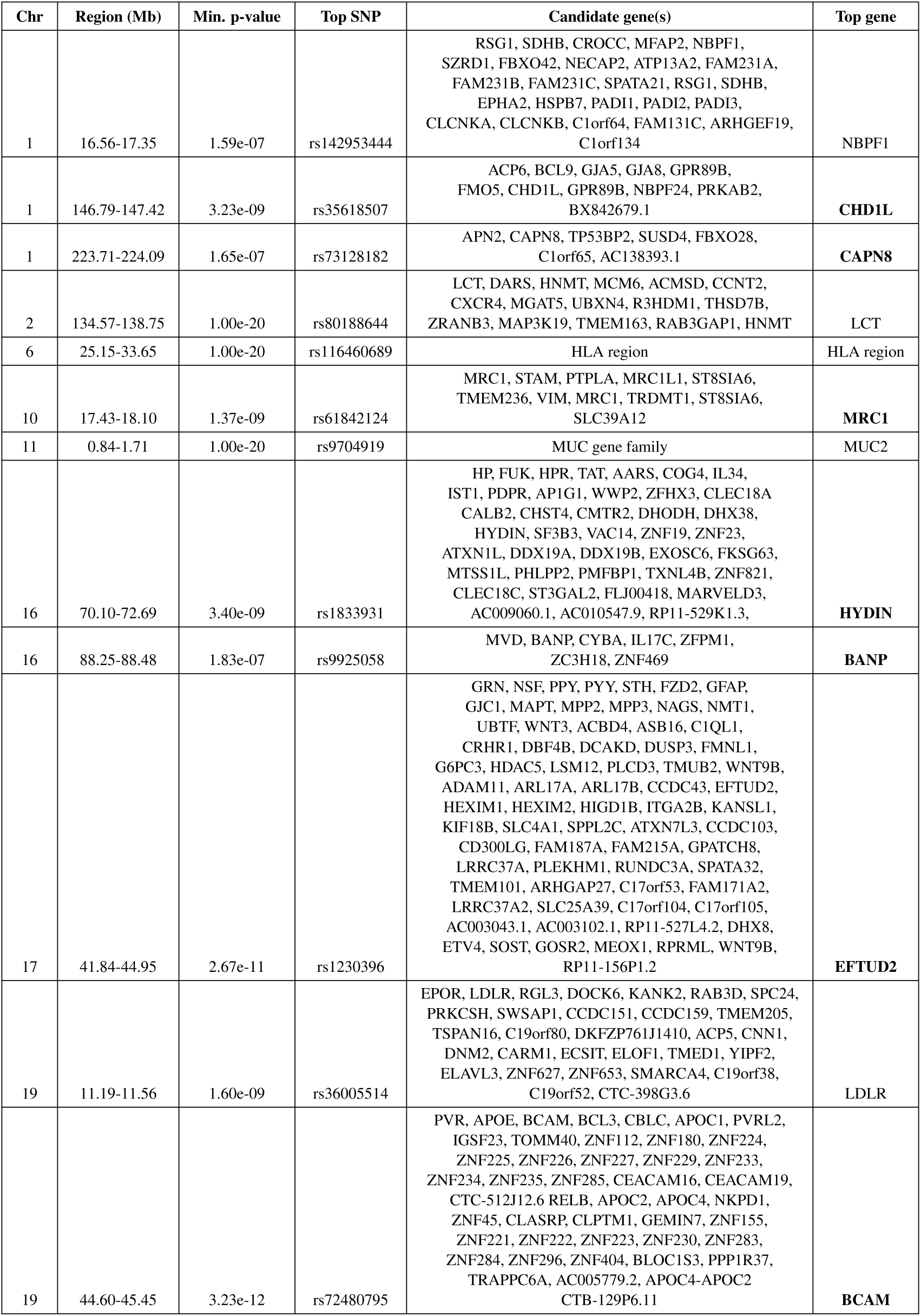
Genome-wide significant selection loci. We report loci with elevated values of the DRC_50_ statistic (in the past 50 generations) in the UKBB (after adjusting for multiple testing p < 0.05*/*52, 003 = 9.6 × 10^−7^). The DRC_50_ statistic of recent positive selection was computed using all 487, 409 individuals from the UKBB. When multiple candidate genes were found, we only retained the one nearest to the top SNP, referred to as the top gene (i.e with the smallest p-value). Novel genes are denoted in bold.

**Supplementary Table 5.** Marker density and recombination rate percentiles for genome-wide significant selection loci under selection. We divided the genome into 0.1 Mb windows and computed recombination rate and marker density within each window. We then ranked the windows by marker density (from high to low values) and by recombination rate (from low to high values) to get percentiles for each window. We associated each of the selection peaks reported in Supplementary Table 4 to the window they fell in. We report the marker density and recombination rate percentiles for each selection peak.

| Chr | Region (Mb) | Min. p-value | Top SNP | Recombination rate at peak | Percentile recombination rate | Marker density at peak | Percentile marker density |
| --- | --- | --- | --- | --- | --- | --- | --- |
| 1 | 16.56-17.35 | 1.59e-07 | rs142953444 | 0.54 | 0.9804 | 12 | 0.9343 |
| 1 | 146.79-147.42 | 3.23e-09 | rs35618507 | 0.17 | 0.687 | 36 | 0.2345 |
| 1 | 223.71-224.09 | 1.65e-07 | rs73128182 | 0.19 | 0.7286 | 39 | 0.1857 |
| 2 | 134.57-138.75 | 1.00e-20 | rs80188644 | 0.1 | 0.5096 | 22 | 0.5945 |
| 6 | 25.15-33.65 | 1.00e-20 | rs116460689 | 0.25 | 0.8159 | 47 | 0.0966 |
| 10 | 17.43-18.10 | 1.37e-09 | rs61842124 | 0.1 | 0.5327 | 8 | 0.9853 |
| 11 | 0.84-1.71 | 1.00e-20 | rs9704919 | 0.3 | 0.8655 | 133 | 0.003 |
| 16 | 70.10-72.69 | 3.40e-09 | rs1833931 | 0.45 | 0.9577 | 19 | 0.7041 |
| 16 | 88.25-88.48 | 1.83e-07 | rs9925058 | 0.19 | 0.7293 | 21 | 0.6344 |
| 17 | 41.84-44.95 | 2.67e-11 | rs1230396 | 0.04 | 0.2594 | 21 | 0.6338 |
| 19 | 11.19-11.56 | 1.60e-09 | rs36005514 | 0.44 | 0.9542 | 577 | 0.0001 |
| 19 | 44.60-45.45 | 3.23e-12 | rs72480795 | 0.45 | 0.9581 | 58 | 0.042 |

**Supplementary Table 6.**
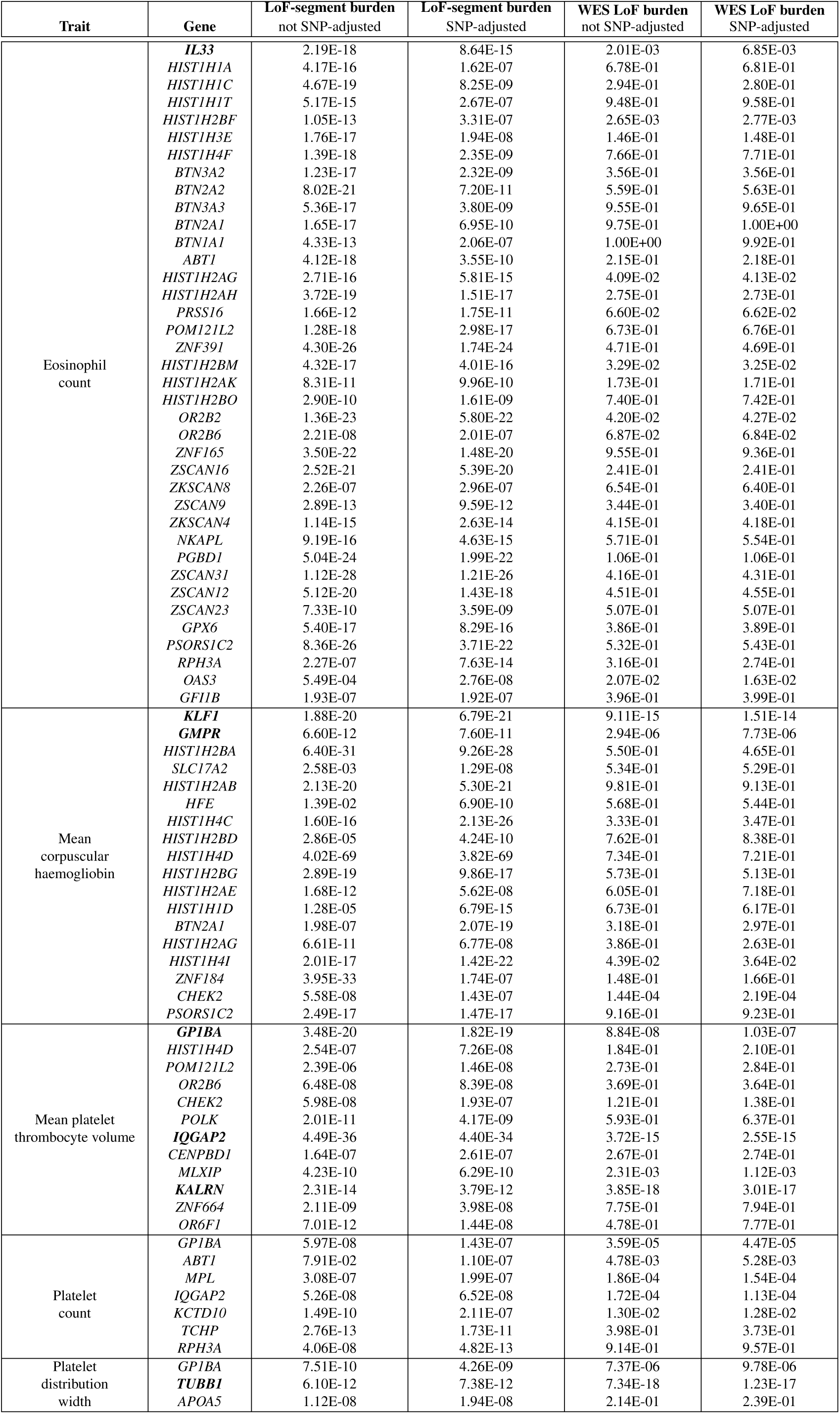
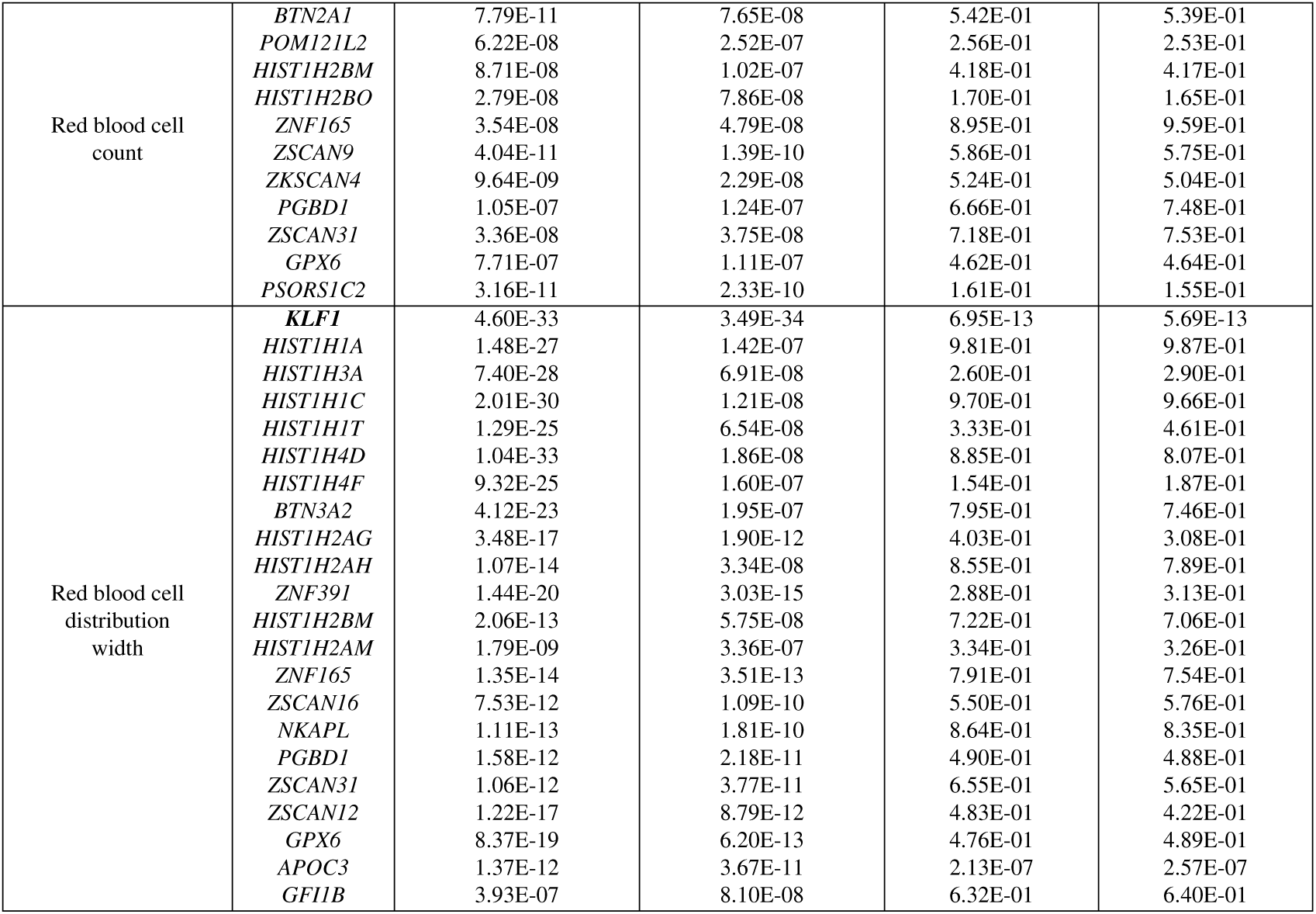
Exome-wide significant genes by LoF-segment burden, and comparison to other tests. A detailed comparison between the four association tests we performed. We report all genes found significant using SNP-adjusted LoF-segment burden (using 303,125 non-sequenced samples; 111 associations in total), as well corresponding p-values for LoF-segment burden without SNP-adjustment, and WES-LoF association (using 34,422 sequenced samples) with or without SNP-adjustment. P-values are computed using *two-sided t-tests*. Genes in bold correspond to hits previously reported by Van Hout et al. (45), for a total of eight replicated signals at exome-wide significance in the non-sequenced cohort. SNP-adjustments resulted in decreased significance for the LoF-segment burden test, but had a smaller effect on the WES-LoF-based approach; in total WES-LoF with SNP-adjustment achieved similar p-values in all but one (*GMPR*) of the exome-wide significant hits obtained by the simple WES-LoF test.

